# Genomic basis of transcriptome dynamics in rice under field conditions

**DOI:** 10.1101/451609

**Authors:** Makoto Kashima, Ryota L. Sakamoto, Hiroki Saito, Satoshi Ohkubo, Ayumi Tezuka, Ayumi Deguchi, Yoichi Hashida, Yuko Kurita, Koji Iwayama, Shunsuke Adachi, Atsushi J. Nagano

**Affiliations:** Research Institute for Food and Agriculture, Ryukoku University, Shiga, Japan; Seibi Senior High School, Gifu, Japan; Graduate School of Agriculture, Kyoto University, Kyoto, Japan; Tropical Agriculture Research Front, Japan International Research Center for Agricultural Sciences, Okinawa, Japan; Faculty of Data Science, Shiga University, Shiga, Japan; Institute of Global Innovation Research, Tokyo University of Agriculture and Technology, Fuchu, Tokyo, Japan; Faculty of Agriculture, Ryukoku University, Shiga, Japan

## Abstract

How genetic variations affect gene expression dynamics of field-grown plants remains unclear. Using statistical analysis of large-scale time-series RNA-sequencing of field-grown rice from chromosome segment substitution lines (CSSLs), we identified 1675 expression dynamics quantitative trait loci (edQTLs) leading to polymorphisms in expression dynamics under field conditions. Based on the edQTL and environmental information, we successfully predicted gene expression under environments different from training environments, and in rice cultivars with more complex genotypes than the CSSLs. Overall, edQTL’ identification helped understanding the genetic architecture of expression dynamics under field conditions, which is difficult to assess with laboratory experiments^1^.The prediction of expression based on edQTL and environmental information will contribute to crop breeding by increasing the accuracy of trait prediction under diverse conditions.

---

Organisms respond to fluctuations in field environments variably depending on their genetic backgrounds, developmental stages, and physiological status. This variability can cause missing heritability in crop breeding and low accuracy in medicine^2–4^. Environmental stimuli can induce transcriptional responses directly and/or indirectly. Measuring transcriptome dynamics is a comprehensive method for assessing environmental responses and their alteration due to genetic, developmental, and physiological factors. The expression quantitative trait loci (eQTL) approach is frequently used to assess the association between genetic variation and gene expression polymorphism^5–9^. Because this approach requires transcriptome data for the complete set of a QTL mapping population under the given conditions^5–9^, only a limited range of environmental conditions are covered. Contrarily, statistical models based on meteorological information, circadian clock, and developmental age have succeeded in describing transcriptome dynamics in the field^10–12^, although these models can only predict the transcriptome of one or few genotypes used as training data^10–12^. Thus, no study has clarified the relationship between genetic variation and transcriptome dynamics under fluctuating field conditions.

Here, we first defined the loci that determine polymorphisms in gene expression dynamics as ‘edQTL’. edQTL can explain polymorphisms in gene expression dynamics under a broad range of environmental conditions, while eQTL can only explain them under the investigated environment (Fig. 1a). Our approach consists of four steps: time-series RNA-sequencing (RNA-Seq), prediction model development, edQTL detection, and evaluation of edQTL (See Methods, Fig. 1b, and Supplementary Fig. 1). We used the rice (*Oryza sativa* L.) cultivars ‘Koshihikari’ (a leading *japonica* cultivar) and ‘Takanari’ (a high-yield *indica* cultivar)^13,14^ that would present substantial polymorphisms in expression dynamics. The 78 reciprocal CSSLs and two backcross inbred lines (BILs) developed from the parental lines were also used^14–16^ (Fig. 1b and Supplementary Fig. 2a). To refelct the effect of plant age (days after seeding) in our model, we prepared four different sets of rice plants transplanted at two-weeks intervals. Sixteen sets (days) of bihourly sampling for 24 h were conducted from May to September 2015 (cropping season in Japan). At each sampling time, the youngest fully expanded leaves were collected from two different genotypes at each transplant set while withered plants were excluded from the sampling, which could not be sampled (Fig. 1c, Supplementary Fig. 2b, and Table S2). The 926 individual leaves were applied for RNA-Seq, to obtain transcriptome data. After filtering samples and genes (Supplementary Fig. 3a, b) and confirming genotypes (Supplementary Fig. 3c and 4), 23,924 expressed genes from 854 samples were used. Correlation analysis of the transcriptome data showed an obvious transcriptome-wide diurnal variation in the expressed genes (Fig. 1d).

**Figure 1.**
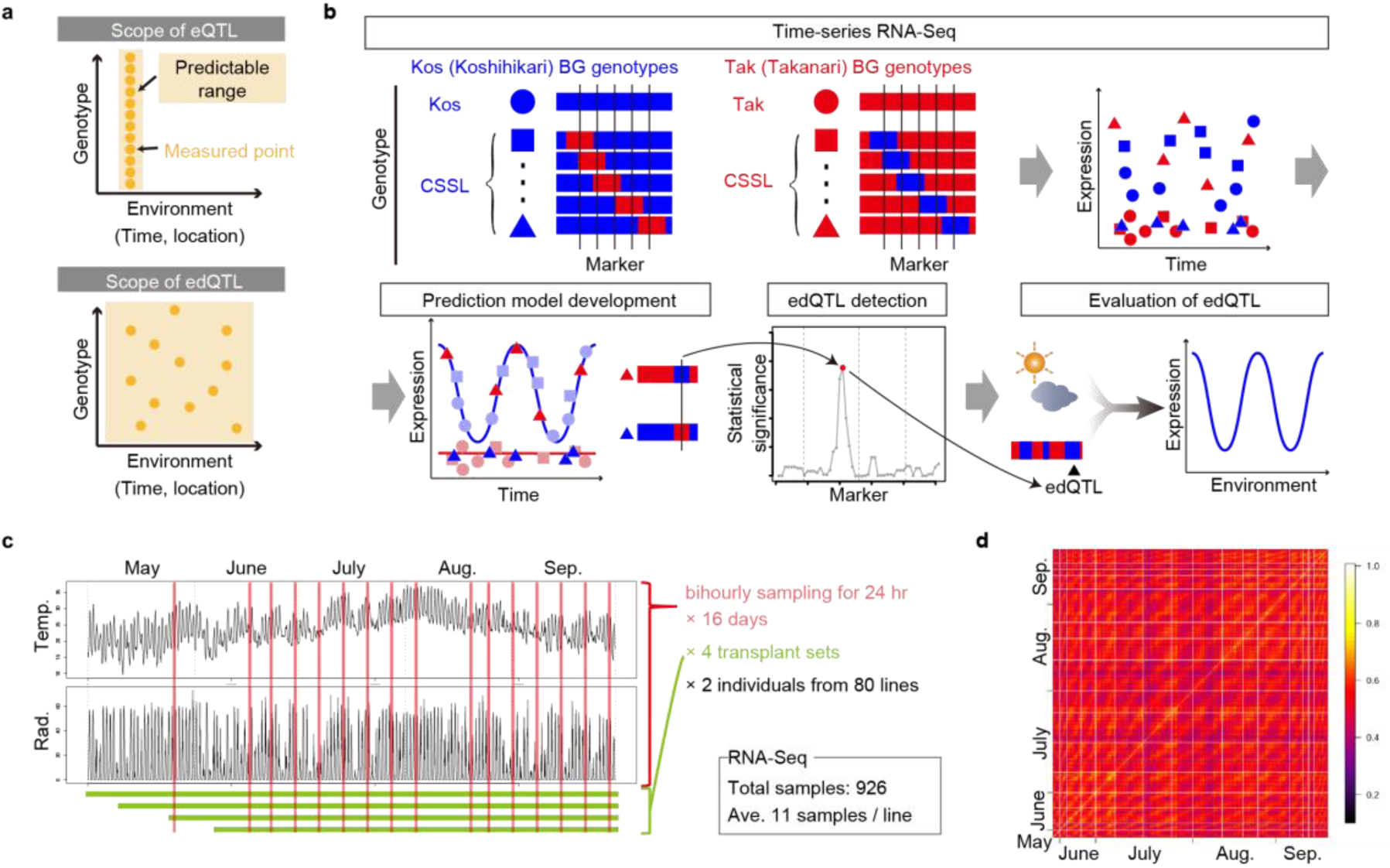
Concept and workflow of expression dynamics quantitative trait loci (edQTL) detection. **a**, Conceptual differences in the scopes of expression quantitative trait loci (eQTL) and edQTL. **b**, Workflow of edQTL detection and its evaluation using chromosome segment substitution lines (CSSLs). **c**, Summary of the sampling design for edQTL detection. The left panels show plots of meteorological data [air temperature (Temp., °C) and global solar radiation (Rad., kJ m^-2^ min^-1^)] in Takatsuki from May to September in 2015. Vertical red lines represent the sampling time points. **d**, Pearson’s correlations among the 854 samples used for developing the prediction model based on expression data of 23,294 expressed genes. White lines indicate the border of each bihourly sampling set.

We then developed two prediction models describing transcriptome dynamics under fluctuating field environments in ‘Koshihikari’ and ‘Takanari’. In this step, we used the statistical modeling tool ‘FIT’^10^ for predicting transcriptome dynamics under field conditions. As most of the genome in the CSSLs was not substituted (Supplementary Fig 2a), the expression dynamics of most genes was expected to be identical to that of the background parent. Thus, the RNA-Seq data of ‘Koshihikari’- and ‘Takanari’-background samples were used to develop the prediction model for ‘Koshihikari’ and ‘Takanari’ lines, respectively. Circadian clock and meteorological data (air temperature and global solar radiation) (Supplementary Fig. 5a) were considered in model development and “scaled age” was used to adjust differences in heading date among rice genotypes (See Methods and Supplementary Fig. 5). We found polymorphisms in the predicted expression dynamics of 3696 genes (15.4% of the expressed genes; Supplementary Fig. 3e and Table S3). For instance, the ‘Koshihikari’ model for *Os09g0343200* showed an obvious diurnal oscillation in expression, whereas the ‘Takanari’-model showed a constant low-level expression (Fig. 2a). Notably, in some ‘Takanari’-background CSSLs, SL1329 and SL1330, the expression of *Os09g0343200* resembled that of the ‘Koshihikari’ model rather than the ‘Takanari’-model (Fig. 2a), suggesting that genetic substitution in the chromosome 9 affected *Os09g0343200* expression.

**Figure 2.**
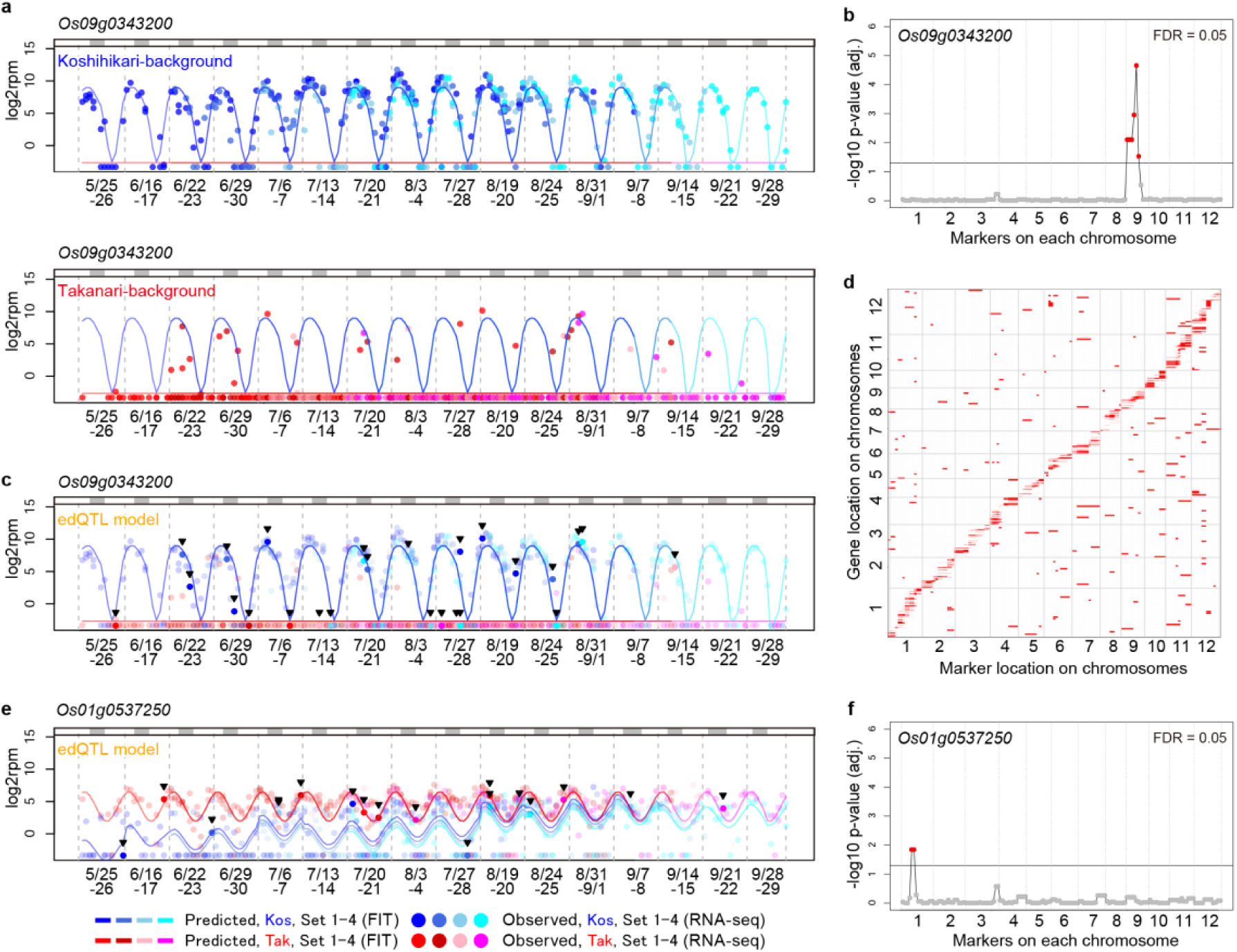
Expression dynamics quantitative trait loci (edQTL) detection revealed *cis*- and *trans*-edQTL that were involved in environmental responses. **a, c, e**, Observed and predicted expressions (log_2_rpm) of *Os09g0343200* (**a**,**c**) and *Os01g0537250* (**e**) in Takatsuki in 2015. The predicted expressions were calculated using ‘FIT’ based on scaled age and environmental information (time, air temperature, and global solar radiation).Blue and red/pink lines indicate predicted expression levels in ‘Koshihikari’ and ‘Takanari’ in transplant sets 1, 2, 3, and 4, respectively. Blue and red/pink points indicate the expression level obtained by RNA-sequencing for samples in transplant sets 1, 2, 3, and 4 of individuals with ‘Koshihikari’-background genotype (BG) and ‘Takanari’-BG in (**a**) or ‘Koshihikari’- and ‘Takanari’-type edQTL in (**c**,**e**), respectively. In (**c**,**e**), strong colored points emphasized by the arrowheads indicate samples harboring edQTL different from their BGs, and light-colored points indicate the samples harboring edQTL identical to their BGs. The upper gray bar indicates dark periods (global solar radiation < 0.3 kJ m^-2^ min^-1^). **b**,**f**, edQTL regulating *Os09g0343200* (**b**) and *Os01g0537250* (**f**). **d**, Position of the 1675 edQTL are shown as red bars (false discovery rate = 0.05). X-axis and Y-axis represent the positions of markers with edQTL and the positions of genes influenced by edQTL, respectively.

The CSSLs had been previously genotyped using 141 simple sequence repeat (SSR) markers (Supplementary Fig. 2a)^14^. As so, at the edQTL detection step, we searched for the SSR markers explaining the expression dynamics polymorphisms. We evaluated the decrease in the residual error of each gene assuming edQTL around each SSR marker (See Methods and Supplementary Fig. 6a). For *Os09g0343200*, the sum of residual errors significantly decreased only by assuming edQTL on chromosome 9 (Fig. 2b and Supplementary Fig. 6b), indicating that this is a *cis*-edQTL. The prediction models chosen for each sample based on edQTL genotypes explained the expression dynamics of *Os09g0343200* better than their background genotypes (Fig. 2c). Overall, edQTL were identified for 1675 genes (45.3% of genes with expression dynamics polymorphism between ‘Koshihikari’ and ‘Takanari’, false discovery rate = 0.05; Supplementary Fig. 6f), including 222 genes affected by *trans*-edQTL (Fig. 2d). Forty-three genes were affected by multiple edQTL. For 33 of these genes, the sum of residual errors based on all edQTL was smaller than that of the most significant edQTL alone (Supplementary Fig. 7). Detailed results for individual genes can be found in our database (https://ps.agr.ryukoku.ac.jp/osa_edqtl). A *cis*-edQTL could explain the expression dynamics polymorphism of *Os01g0537250* that be specifically observed in young plants (Fig. 2e,f). The expression depended on time of day in both parental models, but scaled age was only important in the ‘Koshihikari’ model (Fig. 2e,f and Supplementary Fig. 8). The edQTL-affected 393 genes depending on the time of day and 275 genes depending on scaled age (Supplementary Fig. 9).

In the edQTL evaluation step, we first compared the predicted performances based on the edQTL model and background genotypes model (BG model) under different environments from where training data were obtained for model development. ‘Koshihikari’, ‘Takanari’, and the CSSLs of two transplant sets were cultivated in a field different from that used in 2015 and sampled in August 2016; 139 RNA-Seq datasets were obtained (Supplementary Fig. 2c and Table S2). Environmental factors differed between the 2015 and 2016 fields (Supplementary Fig. 10a). For 91.8% of the 1675 genes affected by the edQTL, the prediction was improved by the edQTL model compared with the BG model (Fig. 3a,b and Supplementary Fig. 10b). Furthermore, the prediction of the expressed genes for the validation dataset showed comparable accuracy to that of the training dataset (Supplementary Fig. 11). Thus, the identified edQTL could explain the expression dynamics polymorphisms in different years and locations. For further verification, two BILs between ‘Koshihikari’ and ‘Takanari’ (HP-a and HP-b)^15,16^ were cultivated at Takatsuki in 2015, and five and six RNA-Seq datasets were obtained for HP-a and HP-b, respectively (Supplementary Fig 2b and Table S2). These lines carried ‘Koshihikari’ alleles at 16.3% and 19.9% of their markers with the genetic background of ‘Takanari’ (Supplementary Fig 2a). To evaluate the performance of the edQTL-based prediction, we calculated the sum of prediction errors of all edQTL-influenced genes using edQTL-, ‘Koshihikari’-, and ‘Takanari’-models. Overall, the edQTL model provided the best prediction (Fig. 3c,d). Permutation analysis of markers in HP-a and HP-b genomes showed the significant advantage of the edQTL model (p < 0.001) (Fig. 3c,d). For instance, as the genotype of the *trans*-edQTL for *Os03g0388300* comprised the ‘Koshihikari’ allele in the genome of HP-a and the ‘Takanari’ allele in the genome of HP-b (Fig. 3e), ‘Koshihikari’- and ‘Takanari’-models were used to predict *Os03g0388300* expression in the edQTL models for HP-a and HP-b, respectively. The expression of *Os03g0388300* fluctuated with time and it was higher in ‘Koshihikari’ than in ‘Takanari’ (Supplementary Fig. 12a). The prediction of *Os03g0388300* expression based on the edQTL models was better than the prediction based on the BG models (Fig. 3f). Regarding *OsKS3* (*Os04g0611700*)^17^ in HP-b, the prediction based on the edQTL model was worse than that based on the BG model (Supplementary Fig. 12b,c). True edQTL might therefore exist around SSR markers 52 or 53, which were not substituted with the ‘Koshihikari’ allele in HP-b (Supplementary Fig. 12b). Such difficulty may occur in some edQTL-based predictions for the BILs, because several edQTLs identified by the CSSLs can be unlinked to the substituted genome regions in the BILs. Finally, we concluded that our approach successfully identified loci linked to expression dynamics polymorphisms in field conditions.

**Figure 3.**
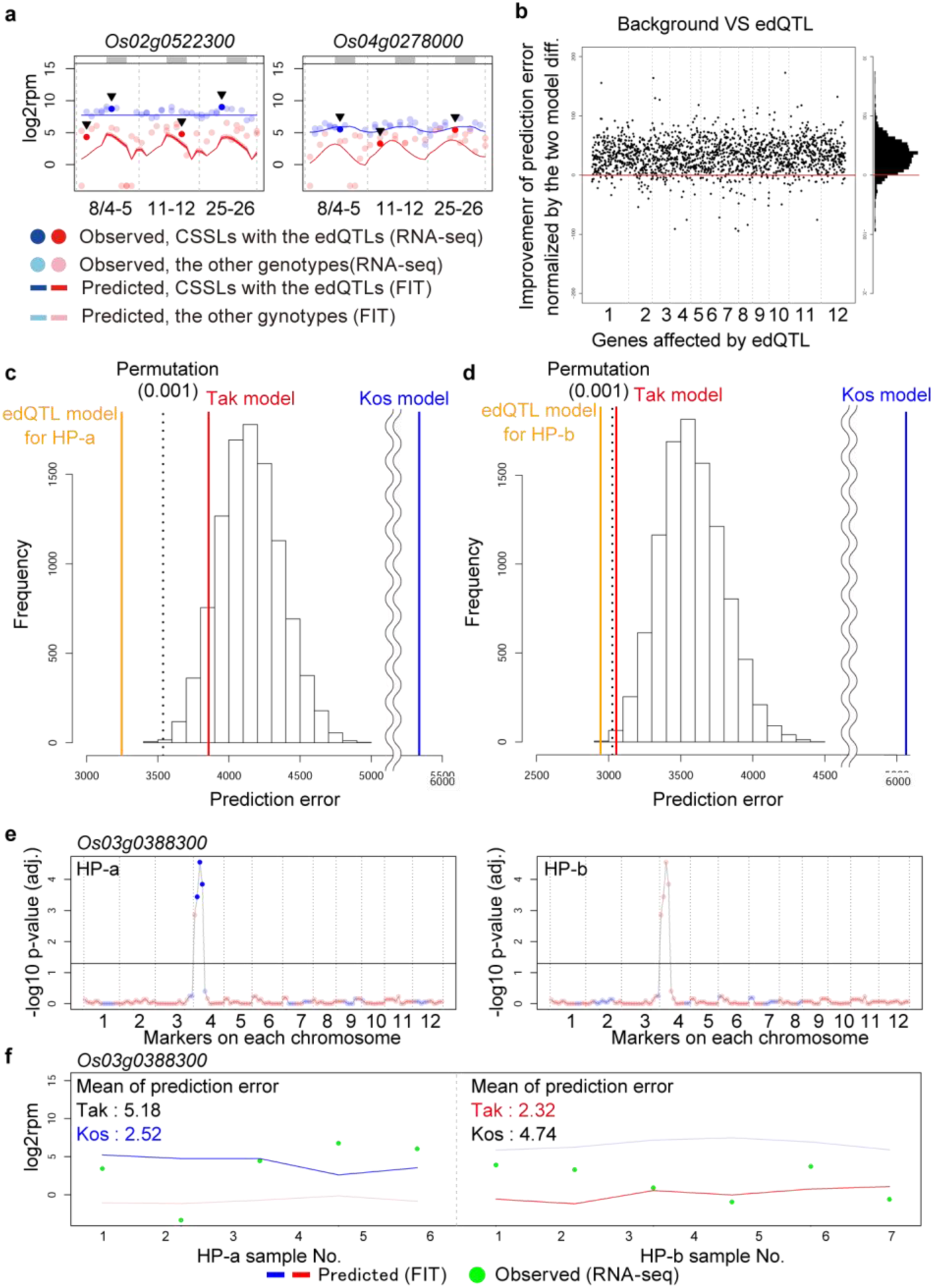
Expression dynamics quantitative trait loci (edQTL)-based prediction of gene expression dynamics under different environments and in cultivars with more complex genotypes than the chromosome segment substitution lines (CSSLs). **a**, Examples of prediction of expression dynamics in Kizugawa in 2016 based on environmental information and edQTL in transplant set 1. Points in intense colors emphasized by arrowheads indicate samples harboring edQTL different from their background genotypes and light colors indicate samples harboring edQTL identical to their background genotypes. The upper gray bars indicate dark periods (global solar radiation < 0.3 kJ m-^2^ min^-1^). **b**, Effects of edQTL over the prediction of gene expression dynamics in Kizugawa in 2016. Genes influenced by edQTL are in order of position on the chromosomes along the horizontal axis. **c**,**d**, Prediction accuracy of edQTL model for HP-a (**c**) and HP-b (**d**). The blue, red, and orange vertical lines indicate the sums of prediction errors based on ‘Koshihikari’, ‘Takanari’, and edQTL models. The histogram shows the distribution of the sums of prediction errors based on the edQTL model in 10,000 permutations of markers in HP-a or HP-b genomes. The dashed vertical line indicates the 0.1% percentile of the distribution. **e**, edQTL for *Os03g0388300* and genotypes of HP-a and HP-b. Dark blue points indicate significant ‘Koshihikari’-type markers. **f**, Prediction of *Os03g0388300* expression in HP-a and HP-b. Intense color lines are applied models for HP-a and HP-b.

The edQTL approach scans a broader range of conditions (Fig. 1a) but it is less sensitive when focusing on a specific condition. This is because eQTL are expected to show the same effects on gene expressions in all samples used in a study. Contrarily, some edQTL were expected to affect gene expressions in some specific samples, depending on their environments. The lower sensitivity of the edQTL approach relative to that of the eQTL approach might explain the smaller fraction of *trans*-edQTL compared with that of *trans*-eQTL in previous studies (13.3% and 62–71%^6,8,9^, respectively) because *trans*-edQTL/eQTL have generally smaller effect sizes than *cis*-edQTL/eQTL^6,8^.

Phenotypic plasticity plays key roles in plant’s environmental adaptation^18,19^. Nevertheless, as this is difficult to study in natural environments, the genetic architecture of phenotypic plasticity under field conditions remains largely unveiled. The edQTL identified by our novel method improve the understanding of the genetic architecture underlying the expression dynamics polymorphisms between ‘Koshihikari’ and ‘Takanari’ in the field. In addition, our method showed that rice gene expression dynamics can be predicted based on genotypic and meteorological information. In crop breeding, polymorphisms of environmental responses result in unexpected performance of a bred crop under environments differing from experimental fields. Because the transcriptome can be beneficial for trait prediction^20–23^ and our approach allows transcriptome predictions under various conditions, it contributes for crop breeding and for understanding plant systems.

## METHODS

### Overview of edQTL identification and verification

Our approach consisted of four steps (Fig. 1b). First, the parent rice lines and their descendants were cultivated in a paddy field and sparsely sampled at several time points for RNA-Seq (Time-series RNA-Seq, Fig. 1b and Supplementary Fig. 1). Second, parental prediction models were developed to describe environmental responses in ‘Koshihikari’ and ‘Takanari’ in terms of gene expression (Prediction model development, Fig. 1b). Most gene expression dynamics would be identical among the CSSLs and the parent with the same background genotype. Thus, we used ‘Koshihikari’- and ‘Takanari’-background CSSLs as well as their parents to develop the parental models (Fig. 1b and Supplementary Fig. 1). Third, the dependency of expression dynamics polymorphisms on genetic variation was statistically evaluated based on comparisons between predictive gene expression and observed gene expression in CSSLs assuming that genetic variation of SSR markers leads to expression dynamics polymorphisms (Supplementary Fig. 6) (edQTL detection, Fig. 1b). Fourth, by integrating the parental models and edQTL information, gene expression dynamics was predicted based on environmental and genotypic information (Evaluation of edQTL).

### Plant materials

We cultivated the following rice (*O. sativa*) lines: the *japonica* variety ‘Koshihikari’, the *indica* variety ‘Takanari’, 78 reciprocal CSSLs (40 ‘Koshihikari’-background lines, except for SL1213, and 38 ‘Takanari’-background lines, except for SL1306)^15^, and two ‘Takanari’-background BILs (HP-a and HP-b)^15,16^. Each variety and line were sown in nursery trays. Approximately one month after sowing, seedlings were transplanted to a paddy field at Takatsuki, Japan (34°51′19″N, 135°37′51″E) in 2015 and at Kizugawa, Japan (34°4′05″N, 135°50′33″E) in 2016. To consider the effect of plant age in the prediction of gene expression dynamics, four and two transplant sets were prepared in 2015 and 2016, respectively (Fig. 1c and Supplementary Fig. 2b,c). Seed sowing and transplanting were conducted according to the following schedules. In 2015, transplant set 1: Seed sowing date “2015-04-03”, Transplanting date “2015-05-01”; transplant set 2: Seed sowing date “2015-04-17”, Transplanting date “2015-05-08”; transplant set 3: Seed sowing date “2015-05-01”, Transplanting date “2015-05-22”; transplant set 4: Seed sowing date “2015-05-15”, Transplanting date “2015-06-05”. In 2016, transplant set 1: Seed sowing date “2016-04-21”, Transplanting date “2016-05-16”; transplant set 2: Seed sowing date “2016-05-19”, Transplanting date “2016-06-08”.

### Sampling and RNA extraction

Sixteen sets (2015) and three sets (2016) of bihourly sampling for 22 h, from 16:00 on one day to 14:00 on the next, were conducted on the following dates (Supplementary Fig. 2b,c and Table S2): 5–6 May, 16–17 June, 22–23 June, 29–30 June, 6–7 July, 13–14 July, 20–21 July, 27–28 July, 3–4 August, 19–20 August, 24–25 August, 31 August to 1 September, 7–8 September, 14–15 September, 21– 22 September, and 28–29 September in 2015; and 4–5, 11–12, and 25–26 August in 2016. We applied a stratified randomization strategy to the sampling schedule to avoid biased sampling of each line over seasons. We separated individual plants into four and two groups in each transplant set in 2015 and 2016, respectively, each containing the same number of individuals per line. Then, the order of sampling was randomized in each group. According to the sampling schedule, two plants from the 82 genotypes of each transplant set were sampled. Because aged rice withered, several samples were missed at the latest cultivation (Supplementary Fig. 2b). Transplant set and sampling time for each sample are listed in Table S2. The youngest fully expanded leaf from each plant was collected, immediately frozen in liquid nitrogen, and stored at -80 °C until RNA isolation for RNA-Seq. Individual plants were only sampled once to avoid wounding response. Thus, all RNA-Seq data were obtained from independent plants. Frozen samples were homogenized with TissueLyser II (Qiagen, Hilden, Germany), and total RNA was then extracted using the Maxwell 16 LEV Plant RNA Kit (Promega, Madison, WI, USA) and Maxwell 16 Automated Purification System (Promega). The concentration of RNA was measured using the Quant-iT RNA Assay Kit, broad range (Thermo Fisher Scientific, Waltham, MA, USA).

### RNA-Seq library preparation and sequencing

An automated liquid handling system Freedom EVO 150 (TECAN, Zurich, Switzerland) and a thermal cycler ODTC 384 (INHECO, Martinsried, Germany) were utilized to prepare the RNA-Seq library of 384 samples simultaneously. The protocol of RNA-Seq library preparation is described below. We enzymatically degraded abundant RNAs such as rRNAs in the leaves, which cause wasteful consumption of sequence reads (Supplementary Fig. 13). Aliquots of 252 and 192 types of 100-μM 60-mer antisense DNAs (IDT, San Jose, CA, USA) covering rRNAs and the top eight most frequently-detected transcripts (*Osp1g00110.1, Osp1g00180.1, Osp1g00420.1, Osp1g00170.1, Osp1g00340.1, Osp1g00330.1, Osp1g00600.1*, and *Os11t0707000-02*) in the RNA-Seq dataset of *O. sativa* leaf were mixed in equal amounts (SDRNA oligo pool)^24,25^. One microgram of total RNA, 1.5 μL of the SDRNA oligo pool, 3 µL of 5× hybridization buffer (0.5 M Tris-HCl (pH 7.4) and 1 M NaCl), and RNase-free water were mixed to a total volume of 10 μL. The mixture was incubated to anneal the antisense DNAs with the target transcripts using the thermal cycler ODTC 384 with the following program: 95 °C for 2 min, slow cooling to 45 °C (0.1 °C/s), and 45 °C for 5 min. Subsequently, 1 μL of Tli Thermostable RNase H (TaKaRa, Kusatsu, Japan), 2 μL of 10× RNase H digestion buffer [500 mM Tris-HCl (pH 7.4), 1 M NaCl, and 200 mM MgCl], and 7 μL of RNase-free water were added and mixed well. This mixture was incubated at 45 °C for 30 min to selectively digest the RNA of the DNA/RNA hybrid double strand. Then, 24 μL of AMPure XP (Beckman Coulter, Brea, CA, USA) beads was added and the mixture was purified in a 384-well magnetic plate following the manufacturer’s instructions. The RNA was then eluted with 14 µL of RNase-free water. Ten microliters of the RNA solution, 1 μL of DNaseI (5 unit/μL), 1 μL of 10× DNaseI buffer (both Promega), 1 μL of 100 mM DTT (Invitrogen, Carlsbad, CA, USA), and 7 μL of RNase-free water were mixed well. This mixture was incubated at 37 °C for 30 min to degrade the SDRNA oligo pool. Subsequently, 24 μL of AMPure XP beads was added and purified according to the manufacturer’s instructions. The RNA was eluted with 14 µL of RNase-free water. Five microliters of the purified mRNA obtained was mixed with 4 μL of 5× SS buffer (Invitrogen), and 1 μL of 100 mM DTT. Fragmentation of mRNA was carried out at 94 °C for 4.5 min and immediately cooled on ice. Subsequently, 0.6 μL of 100 μM random primer (N)_6_ (Promega) and 0.9 μL of RNase-free water were added to the mixture, which was incubated at 50 °C for 5 min and immediately chilled on ice to relax the secondary structure of the RNA. The fragmented RNA with random hexamers and the reverse transcription master mix [1 μL of 100 mM DTT, 0.4 μL of dNTPs (25 mM each; Promega), 0.1 μL of SuperScript IV (Invitrogen), 0.2 μL of actinomycin D (1000 ng/μL) (Nacalai Tesque, Kyoto, Japan), and 5.9 μL of RNase-free water] were mixed. For the reverse transcription step, the mixture was incubated at 25 °C for 10 min, followed by 50 min at 50 °C. SuperScript IV was inactivated by heating the mixture at 75 °C for 15 min. Subsequently, 24 μL of AMPure XP and 12 μL of 99.5% ethanol were added, and the mixture was purified according to the manufacturer’s protocol. The reverse transcription product was eluted with 14 μL of RNase-free water. The purified DNA/RNA hybrid solution without beads and the second-strand synthesis master mix [2 μL of 10× Blue Buffer (Enzymatics, Beverly, MA, USA), 1 μL of dUTP/NTP mix (Fermentas, Burlington, Canada), 0.5 μL of 100 mM DTT, 0.5 μL of RNase H (Enzymatics), 1 μL of DNA polymerase I (Enzymatics), and 5 μL of RNase-free water] were mixed. This mixture was incubated at 16 °C for 4 h, and then purified with 24 μL of AMPure XP according the manufacturer’s protocol. The purified dsDNA was eluted with 14 μL of RNase-free water. Five microliters of the dsDNA solution was used in the following step. End-repair, A-tailing, and adapter ligation were carried out using the KAPA Hyper prep kit (KAPA BIOSYSTEMS, Wilmington, MA, USA) with 1/10× volume of the solutions according the manufacturer’s protocol. One microliter of 0.1 μM Y-shape adapter^24^ was used in the adapter ligation step for 15 min. Then, size selection of the ligation product was performed with 5.5 μL of AMPure XP. The purified dsDNA was eluted with 10 μL of RNase-free water and a second round of size selection was performed with 10 μL of AMPure XP. The size-selected ligation product was eluted with 15 μL of 10 mM Tris-HCl (pH 8.0). One microliter of uracil DNA glycosylase (UDG; Enzymatics) was added to the size-selected ligation product. The mixture was incubated at 37 °C for 30 min to exclude the second-strand DNA. For library amplification, 2 μL of the UDG-digested DNA, 1 μL of 2.5 μM index primer^24^, 1 μL of 10 μM universal primer^24^, 0.5 μL of RNase-free water, and 5 μL of KAPA HiFi HotStart ReadyMix (2×) (KAPA BIOSYSTEMS) were mixed. The DNA fragments with the adapters and an index sequence were amplified using a thermal cycler with the following program: denaturation at 94 °C for 2 min, 16 cycles at 98 °C for 10 s, 65 °C for 30 s, and 72 °C for 30 s for amplification, and 72 °C for 5 min for the final extension. Two rounds of size selection were then performed to remove adapter dimers with equal volume of AMPure XP to the library solution. The purified library was eluted with 10 μL of RNase-free water. The concentration of each library was measured by qPCR using the KAPA SYBR FAST qPCR Master Mix (2×) Kit (KAPA BIOSYSTEMS) and the LightCycler 480 II (Roche Diagnostics, Basel, Switzerland) to mix equal amounts of libraries. One microliter of the mixed library was used for electrophoresis with the Bioanalyzer 2100 and the Agilent High Sensitivity DNA kit (Agilent Technologies, Santa Clara, CA, USA) to check for quality. Sequencing of 50-bp single-ends using HiSeq 2500 (Illumina, San Diego, CA, USA) was carried out by Macrogen (Seoul, South Korea).

### Calculation and normalization of RNA-Seq count data

All obtained reads were processed with Trimmomatic (version 0.33)^26^ using the following parameters: TOPHRED33 ILLUMINACLIP:TruSeq3-SE.fa:2:30:10 LEADING:19 TRAILING:19 SLIDINGWINDOW:30:20 AVGQUAL:20 MINLEN:40. This procedure removed adapter sequences (ILLUMINACLIP:TruSeq3-PE.fa:2:30:10), as well as leading and trailing low quality (Q < 19) or N bases (LEADING:19 TRAILING:19). It also trimmed the reads when the average quality per base dropped below 20 with a 30-base wide sliding window (SLIDINGWINDOW:30:20). Thus, trimmed reads with length > 39 nucleotides and average Q score > 19 were obtained. These trimmed reads were then mapped to the reference sequences of IRGSP-1.0_transcript^27,28^, rRNAs, and transcripts coded in the mitochondria and chloroplast genomes (NC_001320.1, NC_011033.1) using RSEM (version 1.3.0)^29^ and Bowtie (version 1.1.2)^30^ with default parameters. Expected counts of each gene in the RSEM outputs were used in R (version 3.4.2)^31^ for the following analysis. According to a previous study^32^, we removed the effects of misassigned reads in multiplex sequencing experiments with HiSeq 2500 as:

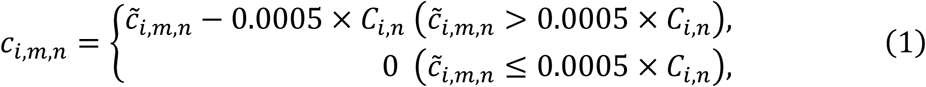

where *c*_*i.m,n*_ and 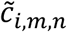 denote corrected and raw read counts of gene *i* in sample *m* in RNA-Seq library pool *n*, respectively. *C*_*i,n*_ denotes the sum of read counts of gene *i* in RNA-Seq library pool *n*.

Then, read counts of gene *i* in sample *m* in different libraries were merged, followed by read count normalization based on total read counts of transcripts except for the antisense-oligo targets (reads per million: rpm):

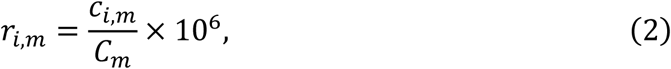

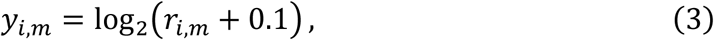

where, *c*_*i,m*_, *r*_*i,m*_, and *y*_*i,m*_ denote merged corrected read counts, normalized read counts, and the log2-transformed value of read counts of gene *i* in sample *m*, respectively. *C*_*m*_ denotes total read counts of transcripts except for the antisense-oligo targets in sample *m*.

Of the 926 samples from 2015 and 143 samples from 2016, we used 887 and 139 RNA-Seq transcriptome datasets, respectively, with more than 10^5^ total read counts for all genes, except for the targets of selective depletion (Supplementary Fig. 3a). We then filtered out rarely detected genes (number of samples with read count > 0; ≤ 20% of all samples in the 2015 dataset) from the following analyses to focus on the 23,924 expressed genes only (Supplementary Fig. 3b).

### Identification of specific ‘Koshihikari’ and ‘Takanari’ single nucleotide polymorphisms

The paired-end reads of ‘Koshihikari’ (SRR1630927) and ‘Takanari’ (DRR018360) genomes were trimmed with Trimmomatic (version 0.33)^26^ using the parameters stated in the previous section, except for TruSeq3-PE.fa for adapter trimming. The reads were then mapped to the reference genome (IRGSP-1.0, plastid: NC_001320.1, mitochondrion: NC_011033.1)^27^ using Bowtie2 (version 2.2.9) with option -N1, which allows a mismatch in seed alignment ^33^. By using SAMtools (version 1.3.1)^34^, the reads mapped at multiple loci and the duplicated reads, except for each representative read, were discarded. Picard (version 2.1.0) (https://broadinstitute.github.io/picard/) was used to add read group information to each SAM file. All SAM files of each sample were merged and a VCF file was generated using SAMtools (version 1.3.1)^34^ and BCFtools (version 1.1)^34^. Single nucleotide polymorphisms (SNPs) were filtered using VCFtools (version 0.1.14)^35^ with option --minDP5, which indicates that all alleles supported by more than four reads are reported. The VCF file generated was analyzed using R (version 3.4.2)^31^ and the R package “vcfR” (version 1.5.0)^36^. We selected 15,117 homozygous SNPs distinguishing ‘Koshihikari’ and ‘Takanari’ genotypes for the following analysis of genotype confirmation.

### Genotype confirmation

In previous studies, the genotypes of the CSSLs and BILs were examined based on the genotypes provided by 141 SSR markers^14^. To confirm the genotype of each sample, we estimated each SSR marker genotype based on the SNPs obtained by RNA-Seq. First, to type ‘Koshihikari’- or ‘Takanari’-type SNPs in each RNA-Seq sample, the trimmed RNA-Seq reads of each sample were mapped to the reference genome as described in the previous section using Bowtie2 (version 2.2.9). Then, the SNPs were filtered using VCFtools (version 0.1.14)^35^ with option --minDP3, which indicates that all alleles supported by more than two reads are reported. To estimate each SSR marker genotype in our samples, ‘Koshihikari’- and ‘Takanari’-type SNPs were counted from the SSR marker position until one of the counts reached “7” (Supplementary Fig. 4a,b). In the cases where the SNPs around an SSR marker position were too sparse, all the SNPs in ±1.5 Mbp from each SSR marker were counted (Supplementary Fig. 4a,c). Then, each estimated SSR marker was categorized into ‘Koshihikari’-type, ‘Takanari’-type, or undetermined based on most of its SNP counts (Supplementary Fig. 4a-c). Any samples of the estimated genotypes not perfectly matching the SL1201, SL1229, or SL1321 genotypes might be due to the low density of SNPs around the substituted SSR markers or to doubtful SNP calls. For the samples with estimated genotypes similar to one of these lines, imputation of estimated SSR marker genotypes was conducted (Supplementary Fig. 4d). Finally, each sample was labeled with the genotype of the most similar SSR marker composition. Thus, 865 samples were confirmed as ‘Koshihikari’, ‘Takanari’, or CSSLs and 11 samples as BILs in 2015, and 139 samples as ‘Koshihikari’, ‘Takanari’, or CSSLs in 2016 (Supplementary Fig. 3d).

### Correlation plot of transcriptomes

Pearson’s correlation coefficients of transcriptome data for all pairwise comparisons of the 854 samples from 2015 were calculated as:

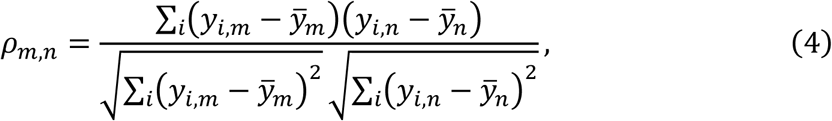

where *ρ*_*m,n*_ denotes the Pearson’s correlation coefficient between samples *m* and *n*. Mean log2-transformed rpms are denoted as 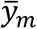 and 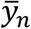, respectively.

The heatmap representing the correlations in the time-series order was drawn with the “image.plot” function in the R package “fields” (version 9.0)^37^.

### Meteorological data

Data of average air temperature per 10 min at the Hirakata Weather Station (34°48′5″N, 135°33′6″E) in 2015 for Takatsuki and at Kyotanabe Weather Station (34°49′8″N, 135°45′6″E) in 2016 for Kizugawa were obtained from the Japan Meteorological Agency. Data of air temperature per minute for the FIT (version 0.0.4)^10^ library were prepared by linear interpolation of the data per 10 min with the “approxfun” function in R (version 3.4.2)^31^. Data of global solar radiation per minute at the Osaka Weather Station (34°40′9″N, 135°37′51″E) in 2015 for Takatsuki and at the Nara Weather Station (34°40′4″N, 135°50′2″E) in 2016 for Kizugawa were also obtained from the Japan Meteorological Agency.

### Conversion of age to scaled age

To consider the difference in plant’s age at heading among genotypes, we defined scaled age. Under the assumption that rice ages linearly before heading at various speeds among genotypes, flowers at scaled age 85, and ages at the same speed after flowering, the scaled age of sample *m* was defined as:

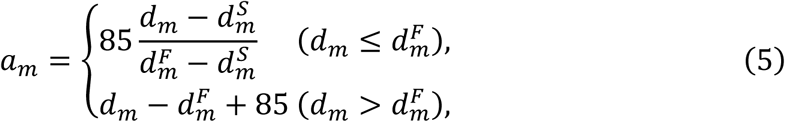

where *d*_*m*_ is the sampling date, 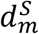 is the seeding date, and 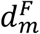 is the flowering date of each line with genotype and seeding date identical to that of sample *m.*

### Calculation of precision weights for each observed normalized read count (log2rpm)

Precision weights were utilized in the training step of FIT (version 0.0.4)^10^ to deal with over-dispersion of RNA-Seq data^10^. Precision weight for each observed log2rpm value was calculated based on the residuals from the smoothed time-series as in the voom method^10,38^ (Supplementary Fig. 14). First, for each gene, spline interpolation against time and observed log2rpm was conducted with the “splinefun” function in R, to infer the function value *s*_*iG*_(*t*) for interpolation of observed log2rpm of gene *i* at time *t* in *G* ∈ {K, T}, where K and T denote ‘Koshihikari’ and ‘Takanari’-background genotypes, respectively (Supplementary Fig. 14a). Then, residual standard deviation of gene *i* in *G*-background genotype *σ*_*i,G*_ was calculated as:

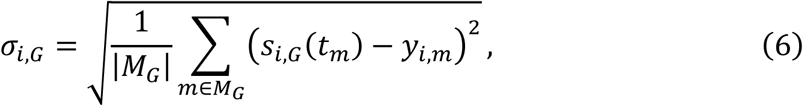

where *t*_*m*_ denotes sampling time of sample *m. M*_*G*_ and |*M*_*G*_| denote indices of samples whose background genotypes are *G* and its size, respectively.

The mean log2-transformed read count of gene *i* in *G*-background genotype was then calculated as:

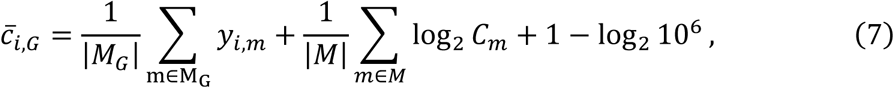

where *M* denotes sample indices in both background genotypes.

A LOWESS curve^39^ was then fitted to the square-root of residual standard deviations as a function of mean log2-transformed read counts, resulting in *l*(), which represents the trend between log2-transformed read counts and square-root of residual standard deviations (Supplementary Fig. 14b).

Similar to 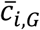, smoothed log2rpm of gene *i* in sample *m* was converted to smoothed log2-transformed read counts of gene *i* in sample *m* as:

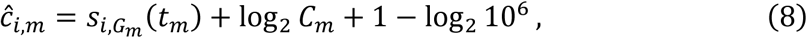

where *G*_*m*_ denotes the background genotype of sample *m*.

Finally, precision weight for observed log2rpm of gene *i* in sample *m* was calculated as (Supplementary Fig. 14c):

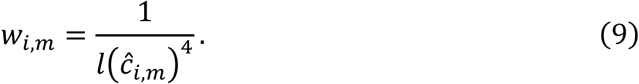

### Prediction model development for ‘Koshihikari’ and ‘Takanari’

A prediction model for gene expression dynamics in ‘Koshihikari’ and ‘Takanari’ was developed with the R package “FIT” (version 0.0.4)^10^. In this section, we describe how “FIT” developed a prediction model.

There are several strategies for the integrative analysis of the transcriptome and meteorological data to investigate environmental responses in field conditions^40–44^. In the FIT package, we employed the statistical modeling approach for a comprehensive relationship among these data and for understanding the effects of age and circadian clock of plants. We used the transcriptome data in combination with the meteorological data, namely air temperature and global solar radiation, which were measured at a meteorological station near the field from where the samples were acquired. Because there was no remarkable disease or pest damage during cultivation and no difference in fertilization among samples, the model did not include these effects.

The log2rpm of the normalized expression for gene *i* of sample *m* was described by a simple linear model:

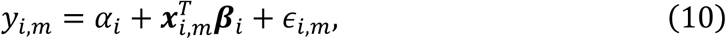

where α_i_, ***x***_*i,m*_, and ***β***_*i*_ are a constant term, a vector of eight explanatory variables, and regression coefficients, respectively. The third term *ε*_*i,m*_ is the independently and identically distributed noise drawn from a Gaussian distribution. The explanatory variables were plant’s scaled age, circadian clock, optional variable, response to environmental stimuli, and the interactions between scaled age and circadian clock and between scaled age and response to environmental stimuli. The optional variable was not used in this study. The plant’s scaled age corresponded to the scaled number of days after transplanting. See section “Conversion of age to scaled age” for the definition of scaled age. The values of the scaled age were normalized to mean 0 and variance 1. The circadian clock, with the peak at an arbitrary time, was represented by two variables corresponding to its cosine and sine components:

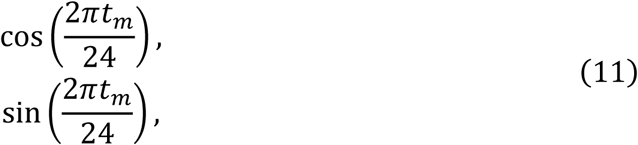

where *t*_*m*_ is the time at which sample *m* was obtained. Similarly, the interaction between scaled age and circadian clock of a plant was represented by two variables.

Although we employed a linear model for simplicity, the actual gene expression can exhibit more complex responses. The complexity of fluctuations in gene expression is mainly explained by the nonlinearity of the response to the environmental stimuli, which is the cumulative sum of the nonlinearly transformed environmental stimuli for a given period. Such cumulative sum of temperature, for example, is designated as cumulative temperature and it is widely used for the prediction of reproduction timing and vegetation in ecology and of harvest timing in agriculture^45,46^. By applying this to gene expression, the environmental response of gene *i* in sample *m* can be described as:

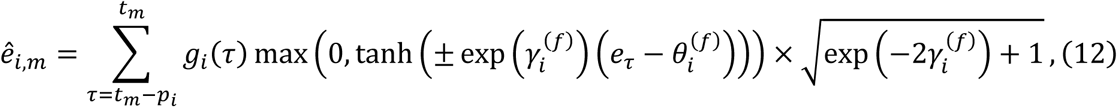

where *p*_*i*_ is the period during which gene expression was affected by an environmental stimulus, *e*_*τ*_ is the value of a meteorological parameter at time *τ* normalized to mean 0 and variance 1, 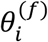 is the response threshold to the stimulus, and 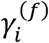 is the parameter that controls the shape of the response (Supplementary Fig. 15). Here, the meteorological data consisted of air temperature and global solar radiation. Of these parameters, the one yielding the highest likelihood was selected for each gene in the optimization procedure. In the limit 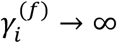, a gene responds to an environmental stimulus in a dose-independent manner, that is, the response is constant if a meteorological parameter exceeds the threshold (Supplementary Fig. 15). Conversely, as this parameter approaches minus infinity, the response also approaches the dose-dependent manner (Supplementary Fig. 15). The sign of the value in the hyperbolic function determines whether the gene responds to an environmental stimulus larger than (positive) or smaller than (negative) the threshold. This sign was decided in the optimization procedure based on the parameter likelihood. The last term 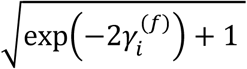 ensures that the scale of the response is largely independent of this parameter. The function *g*_*i*_(*τ*) is a gate function that explains the time-of-day-specific environmental responses. A gate function is defined as:

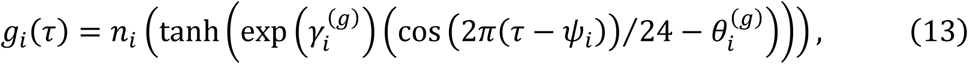

where 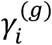 controls the shape of the gate as 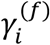 in the environmental response, 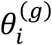 determines the opening length of the gate, and *n*_*i*_() is the function which normalizes an input value [0,1]. This gate function approaches a sinusoidal and rectangular wave with the small and large values of 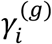, respectively. If the value of 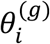 is smaller than −1, the gate is a constant value in the limit 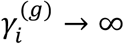, that is, the gate is always open (Supplementary Fig. 15).

It can be considered that some explanatory variables might contribute to neither the explanation nor the prediction of the expression of many genes. Thus, we simultaneously preformed the optimization and variable selection of the regression coefficients by using group lasso^47^. Let *I* be the index set for scaled age, environmental response, and the interaction between scaled age and environmental response. The cost function to be minimized is thus defined as:

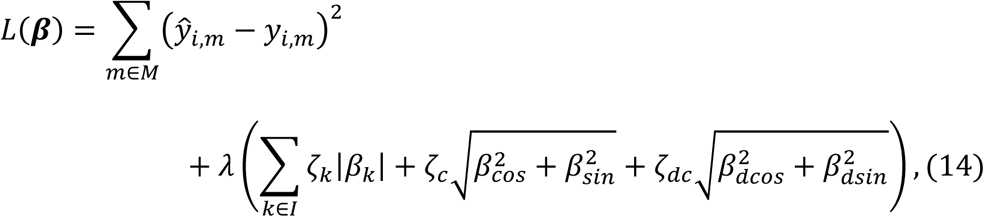

where *λ* and ***ζ*** are the regularization parameter and the adaptive weights, respectively.

The values of the adaptive weights are decided as in the adaptive lasso^48^ or the adaptive group lasso^49^. Further, the adaptive weights were used to absorb differences in the degrees of freedom by multiplying the covariates of the environmental response and interaction between the environmental response and scaled age by seven, as seven free parameters were considered. We selected the value of *λ* as the largest value for which the cross validation (CV) error was smaller than the sum of the minimum CV error and its standard error.

The values of the parameters related to environmental responses, which are 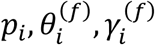, and the sign of the value in the hyperbolic function in Eq. (12), and 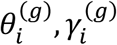, and *ψ*_*i*_; in Eq. (13), are selected by the Nelder-Mead algorithm^50^. Because the likelihood function is complex and has multiple local maxima, a grid search was performed before optimization. In each step of the optimization by the Nelder-Mead algorithm, we performed an ordinary least square regression instead of group lasso (Eq. (14)) to reduce the computational cost.

‘Koshihikari’ and ‘Takanari’-prediction models were developed based on time-series RNA-Seq data of the ‘Koshihikari’-background CSSLs and ‘Koshihikari’ and of the ‘Takanari’-background CSSLs and ‘Takanari’ in 2015, respectively (Supplementary Fig. 2b). Values of the expressed genes (a matrix of log2rpm for each sample), sample attributes (a matrix of sampling year, month, day, hour, and min and scaled age for each sample), weather data (a matrix of air temperature and global solar radiation at every minute in 2015), and precision weights for the matrix of log2rpm of each sample (Supplementary Fig. 16) were prepared. Details on the parameters for ‘FIT’ are listed in https://github.com/naganolab/Rice_edQTL-analysis_and_edQTL-based-prediction/analysis.R. As a result, the functions *f*_*i,G*_() were yielded for gene *i*. A log2rpm of gene *i* of sample *m* was predicted as (Supplementary Fig. 16):

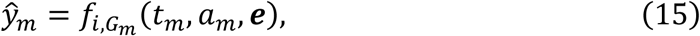

where ***e*** denotes meteorological data (air temperature and global solar radiation) at every minute.

### Detection of genes with expression dynamics polymorphism

To increase detection power in multiple tests, we focused on genes with expression dynamics polymorphism. Euclidean distances of gene *i* between ‘Koshihikari’- and ‘Takanari’-prediction models (*d*_*i*_) were calculated as:

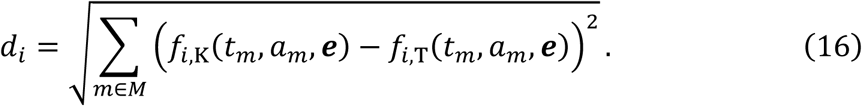

As a result, 3696 genes with *d*_*i*_ > 40 were defined as genes with expression dynamics polymorphism (Supplementary Fig. 3e).

### Detection of edQTL

According to background genotypes, the sum of residual errors between the observed log2rpm and predicted log2rpm of gene *i* in the ‘Koshihikari’- or ‘Takanari’-models (*E*_*i*_) were calculated as (Supplementary Fig. 3a):

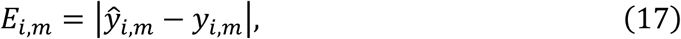

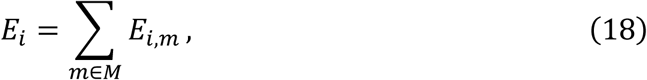

where *E*_*im*_ denotes the residual error between the observed log2rpm and predicted log2rpm of gene *i* in sample *m* on the assumption that the ‘Koshihikari’- or ‘Takanari’-type gene expression dynamics shown by sample *m* is determined based on background genotype *G*_*m*_. Similarly, on the assumption that edQTL for gene *i* exist around SSR marker *g*, the sum of residual errors between the observed log2rpm and predicted log2rpm of gene *i* in the ‘Koshihikari’- or ‘Takanari’-models 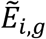 were calculated as (Supplementary Fig. 3a):

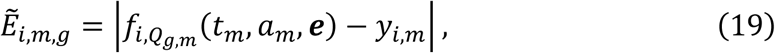

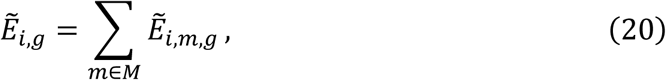

where 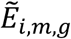 denotes the residual error between the observed log2rpm and predicted log2rpm of gene *i* in sample *m* and *Q*_*g,m*_ denotes genotype of SSR marker *g* in sample *m*. Thus, 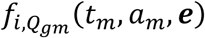 denotes predicted log2rpm in sample *m* assuming the existence of edQTL for gene *i* around SSR marker *g*. The improvement in residual errors assuming that edQTL for gene *i* exists around *g* was therefore calculated as (Supplementary Fig. 3a,b):

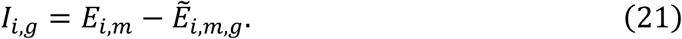

To evaluate the statistical significance of the improvement in the sum of residual errors, 1000 permutations of samples with ‘Koshihikari’- and ‘Takanari’ genetic background were performed (Supplementary Fig. 6e). For each permutation, the improvement in residual errors on the assumption that edQTL for gene *i* exist around SSR marker *g* was calculated (Supplementary Fig. 6b). Then, we generated the null distribution of improvements in residual errors by fitting log-normal distribution to obtained improvements using the fitdist function in the R package “fitdistrplus”^51^, with the fitting method of quantile matching of 70% and 1% (Supplementary Fig. 6c). For almost all genes with expression dynamics polymorphism, the 99.9 percentiles of the fitted log-normal distributions were higher than (2863/3696 genes) or almost the same as those of the distribution obtained by the permutations (Supplementary Fig. 6d), indicating that the *p*-value obtained from log-normal distribution was conservative. Thus, the *p*-value for the improvement in the sum of residual errors for gene *i* on the assumption that an edQTL affecting gene expression dynamics of gene *i* exists around SSR marker *g* was obtained based on the null distribution.

The adjustment for multiple comparisons against *p*-values was performed with the Benjamini-Hochberg method^52^ using the p.adjust function in R. Finally, we defined the markers with the smallest adjusted *p*-value (< 0.05) in each peak for each gene as the edQTL affecting each gene expression dynamics.

### edQTL-based prediction of gene expression dynamics (edQTL model)

To predict gene expression with edQTL information, the ‘Koshihikari’- or ‘Takanari’-model was selected based on the genotype of the edQTL for edQTL-affected genes and of genetic backgrounds for the other expressed genes. Let 𝒢_i_ denote the set of edQTLs affecting gene *i*. We selected the model for prediction of log2rpm of gene *i* in sample *m* as:

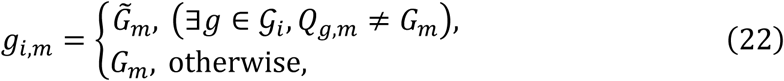

where 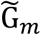 denotes the opposite genotype to that of the genetic background in sample *m*, that is, 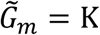, if sample *m* is ‘Takanari’-background, and vice versa. Then, predicted log2rpm of gene *i* in sample *m* was calculated as:

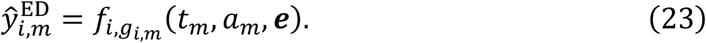

### Statistical test for the accuracy of gene expression dynamics prediction in BILs

The sums of prediction errors of all genes affected by each edQTL for the BILs based on the ‘Koshihikari’ model (*E*_K_), ‘Takanari’ model (*E*_T_), and edQTL model (*E*^ED^) were calculated as:

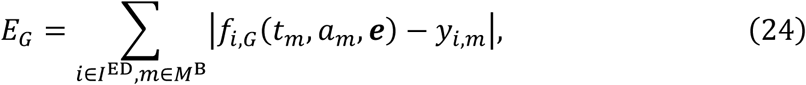

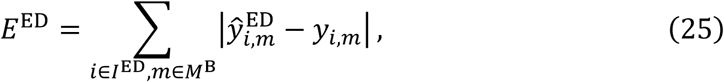

where *M*^B^ denotes the indices of BIL samples and *I*^ED^ denotes the genes affected by the edQTL.

To evaluate the statistical significance of improvement in the sums of prediction errors, 10,000 permutations were performed for the markers in each BIL genome. In each permutation, the sums of prediction errors of all genes affected by the edQTL for the BILs were calculated with the edQTL model. Then, the value corresponding to percentile 0.1 of the sums of prediction errors was calculated.

## Code availability

The scripts used in this study are available from https://github.com/naganolab/Rice_edQTL-analysis_and_edQTL-based-prediction.

## Data availability

All datasets generated and/or used in this study are available from PRJDB7234.

## Supplementary Information

is available in the online version of the paper.

## Acknowledgments

We thank the field work support team for help at field sites, Fumie Kobayashi for assistance with laboratory experiments, and Yasuhiro Sato for valuable discussions. We thank Toshio Yamamoto (National Agriculture and Food Research Organization) for providing seeds of the CSSLs and BILs between ‘Koshihikari’ and ‘Takanari’. This work was supported by the JST CREST JPMJCR15O2 and JSPS KAKENHI Grants JP16H06171 and JP16H01473 to A.J.N.

## Author contributions

M.K. analyzed the data. R.L.S., H.S., S.O. and S. A. performed field work. M.K., A.T., A.D., Y.H., and Y.K. conducted RNA-Seq experiments. K.I. developed the FIT library. A.J.N. designed the study. M.K. and A.J.N. wrote the manuscript with input from all co-authors.

## Supplemental Tables

**Table S1 SSR marker composition in the rice lines used in the present study**

“Position” indicates the position of each SSR marker on the chromosomes in the IRGSP1.0 reference genome. A and B indicate ‘Koshihikari’- and ‘Takanari’-type alleles, respectively.

**Table S2 Sample attributes used in this study**

“TransPlantSet” indicates the transplant sets in each year. Details of seeding and transplant dates in each set are described in the methods. “Year”, “month”, “day”, and “hour” columns indicate the sampling date. The “LineName” column indicates the predicted genotypes based on SNPs detected in each RNA-Seq. The “field” column indicates the location of the fields.

**Table S3 *Oryza sativa* genes and their expressions**

The “Expression” column indicates whether the genes are “expressed genes” or not. “Yes” indicates that “expressed genes” were detected in more than 20% of all samples. The “Polymorphism” column indicates whether the genes show expression dynamics polymorphism between ‘Koshihikari’ and ‘Takanari’ or not. “Yes” indicates that the Euclidian distances of predicted expression dynamics between ‘Koshihikari’ and ‘Takanari’ were more than 40. The “edQTL”, “cis.edQTL”, and “trans.edQTL” columns indicate whether the genes have the designated edQTL or not.

