## supplemental_data for "Genomic basis of transcriptome dynamics in rice under field conditions"

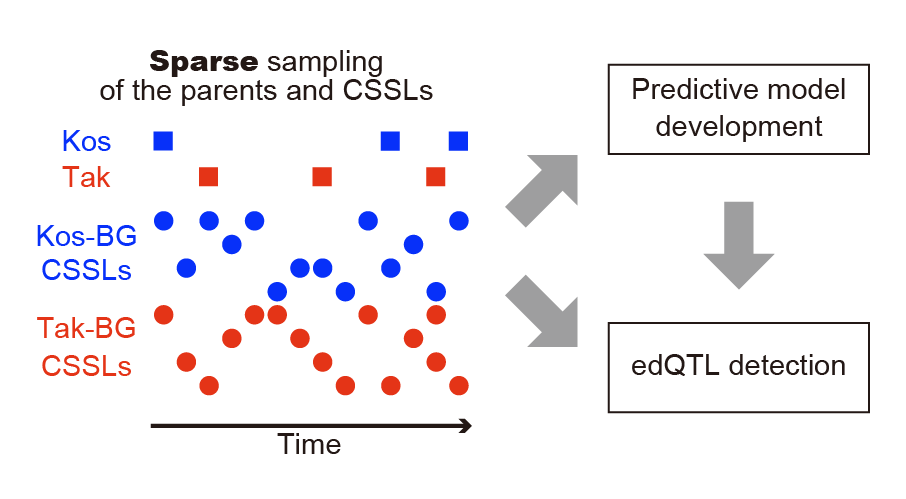

**Supplementary Figure 1 | Strategy for predictive model development and edQTL detection.** Illustration of the strategy for edQTL detection in this study. The sparsely sampled parents, ‘Koshihikari’ (Kos) and ‘Takanari’ (Tak), and Kos-background-genotype (BG) chromosome segment substitution lines (Kos-BG CSSLs) and Tak-BG CSSLs were used in both model development and edQTL detection.

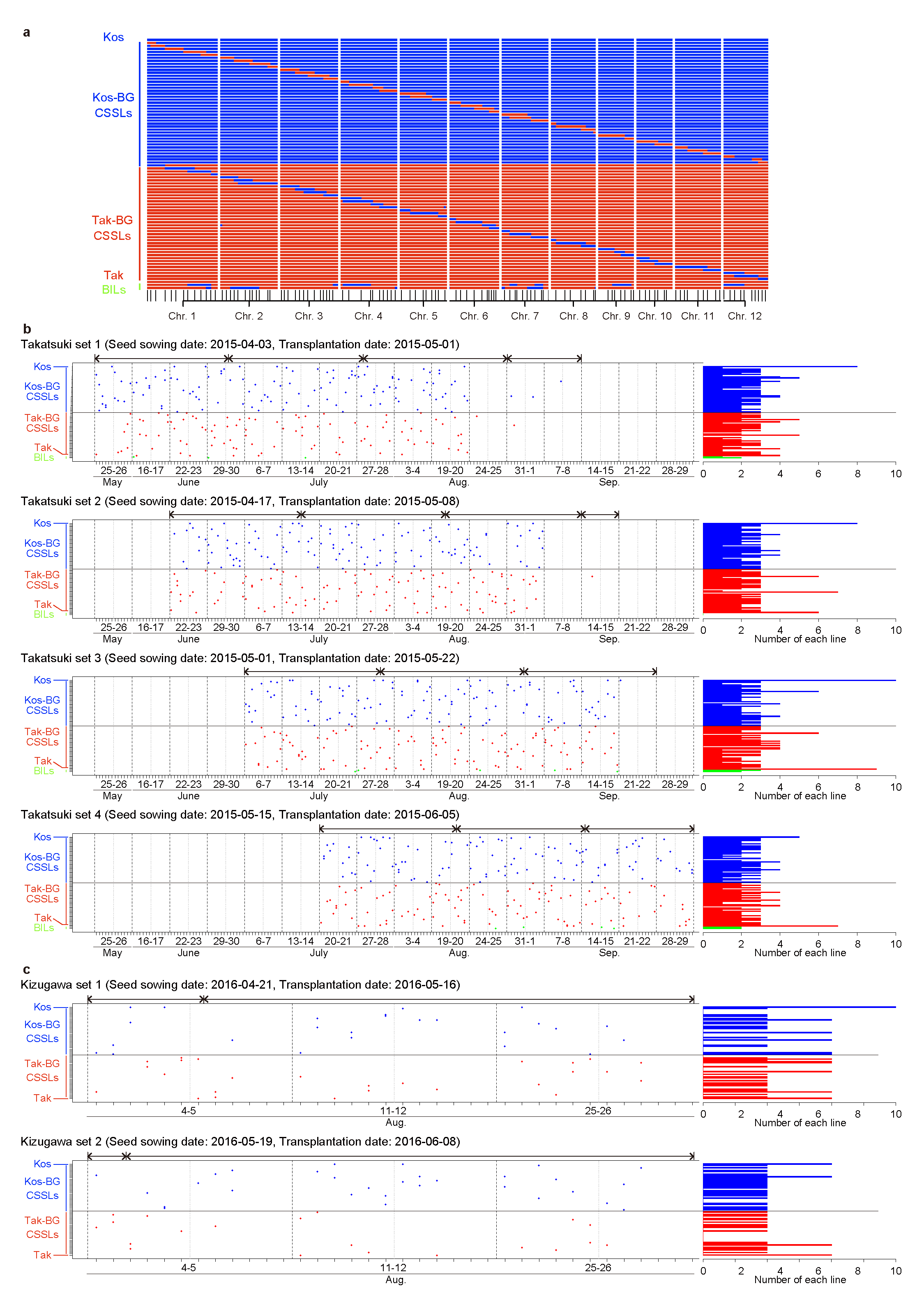

**Supplementary Figure 2 | Samples used in this study. a,** Graphical genotypes of the parental lines, 78 reciprocal chromosome segment substitution lines (CSSLs), and two backcross inbred lines (BILs). Blue and red regions indicate chromosome segments derived from ‘Koshihikari’ and ‘Takanari’, respectively. The lower black vertical lines indicate the positions of the SSR markers. **b,c,** A summary of bihourly sampling for 22 h in this study. Each sampling was conducted from 16:00 to 14:00. Each point indicates a sampled line. The histograms show the total number of samples of each line from each transplant set. The upper double-headed arrows indicate groups for the stratified randomization. Sample details are described in Table S2.

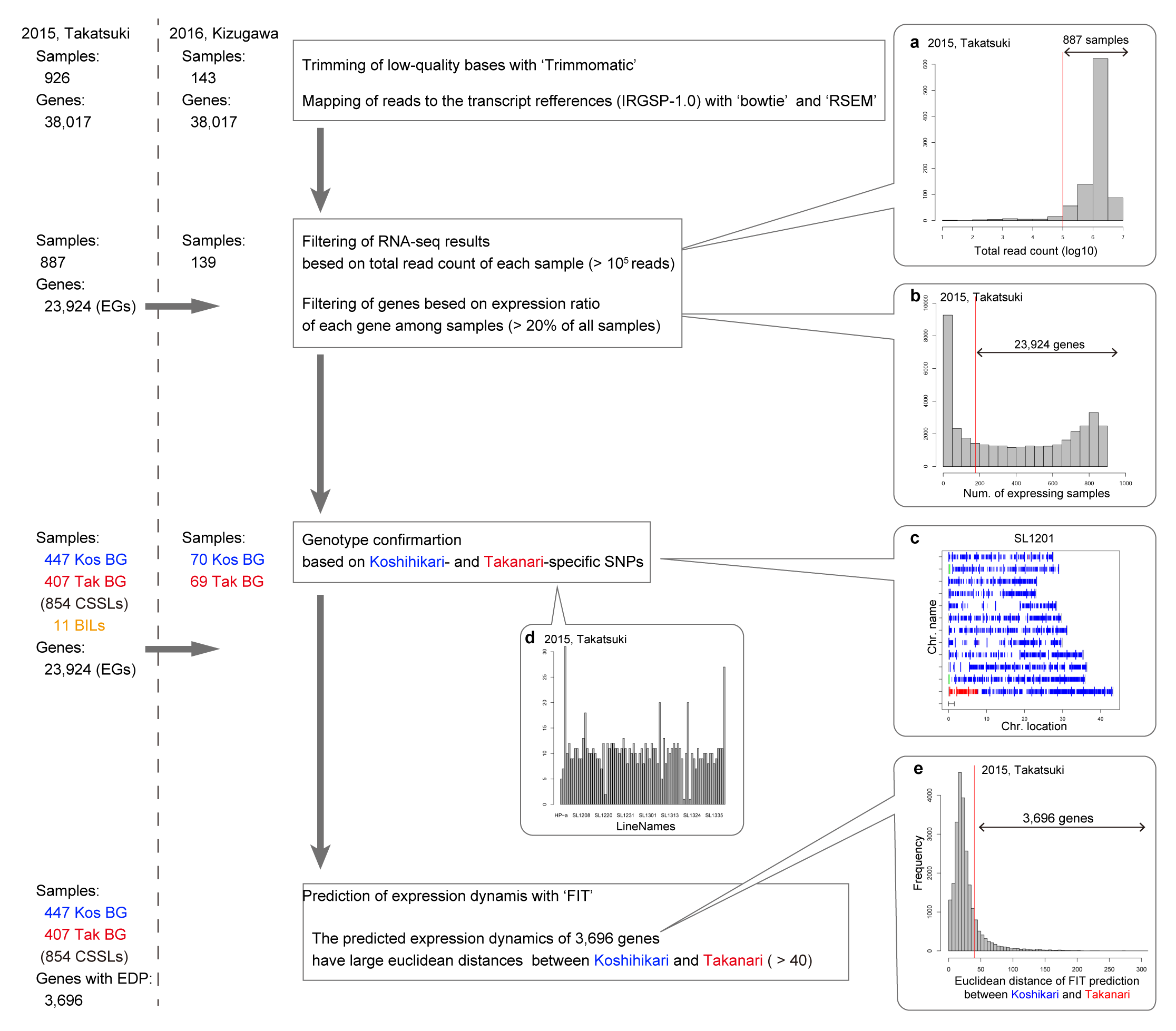

**Supplementary Figure 3 | Workflow of the RNA-Seq data preprocessing. a,** Total read counts of RNA-Seq data in the 2015 dataset, except for selective-depletion target genes. Among the 926 samples, 887 had more than 10^5^ total read counts. **b,** Number of detected samples of each gene in the 2015 dataset. Among the 38,017 genes, 23,924 expressed genes (EGs) were detected in more than 20% of the 887 samples. **c,** Map of the single nucleotide polymorphisms (SNPs) that can identify chromosome segments of ‘Koshihikari’ and ‘Takanari’. The shorter horizontal lines indicate each SNP and the longer lines indicate each SSR marker. Blue indicates ‘Koshihikari’-type SNP or SSR makers and red indicates ‘Takanari’-type SNP or SSR markers. Green long vertical lines indicate untyped SSR markers. **d,** Summary of the total number of samples of each line in the 2015 dataset. **e,** Euclidean distance of each gene between Koshihikari and Takanari based on expression levels predicted by ‘FIT’ in the 2015 data set. The 3,696 genes showed Euclidean distance > 40 and therefore were defined as genes with expression dynamics polymorphism (EDP).

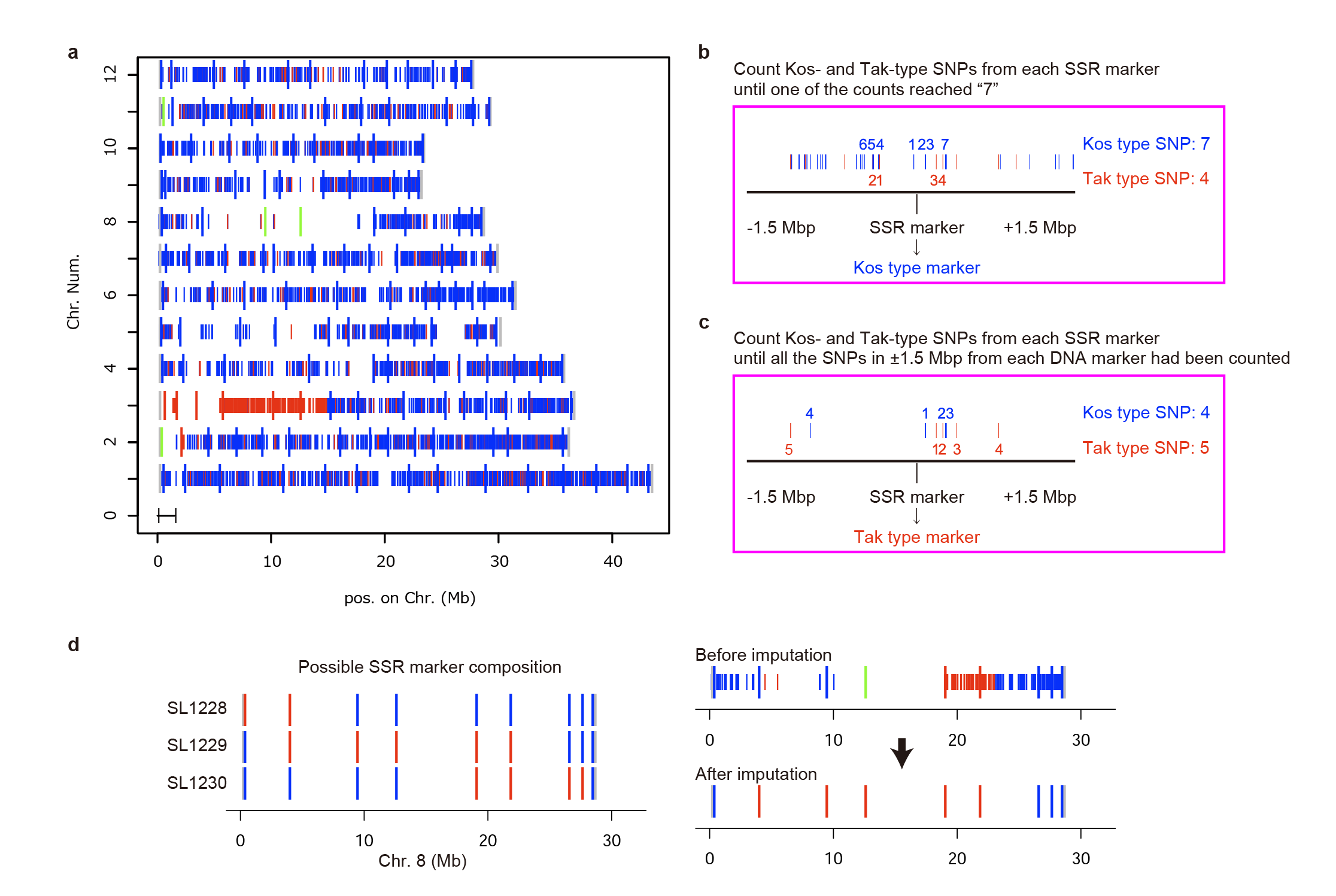

**Supplementary Figure 4 | Schematic illustration of genotype confirmation of CSSLs using RNA-Seq data. a,** Example of genotype confirmation using RNA-Seq data. The shorter horizontal lines indicate each SNP and the longer lines indicate each SSR marker. Blue indicates ‘Koshihikari’-type SNP or estimated SSR makers and red indicates ‘Takanari’-type markers or estimated SSR markers. Green long vertical lines indicate SSR markers that could not be estimated. **b,** Example of SSR marker genotype estimation with dense SNPs. **c,** Example of SSR marker genotype estimation with sparse SNPs. **d,** Example of SSR marker genotype imputation for samples most similar to SL1229.

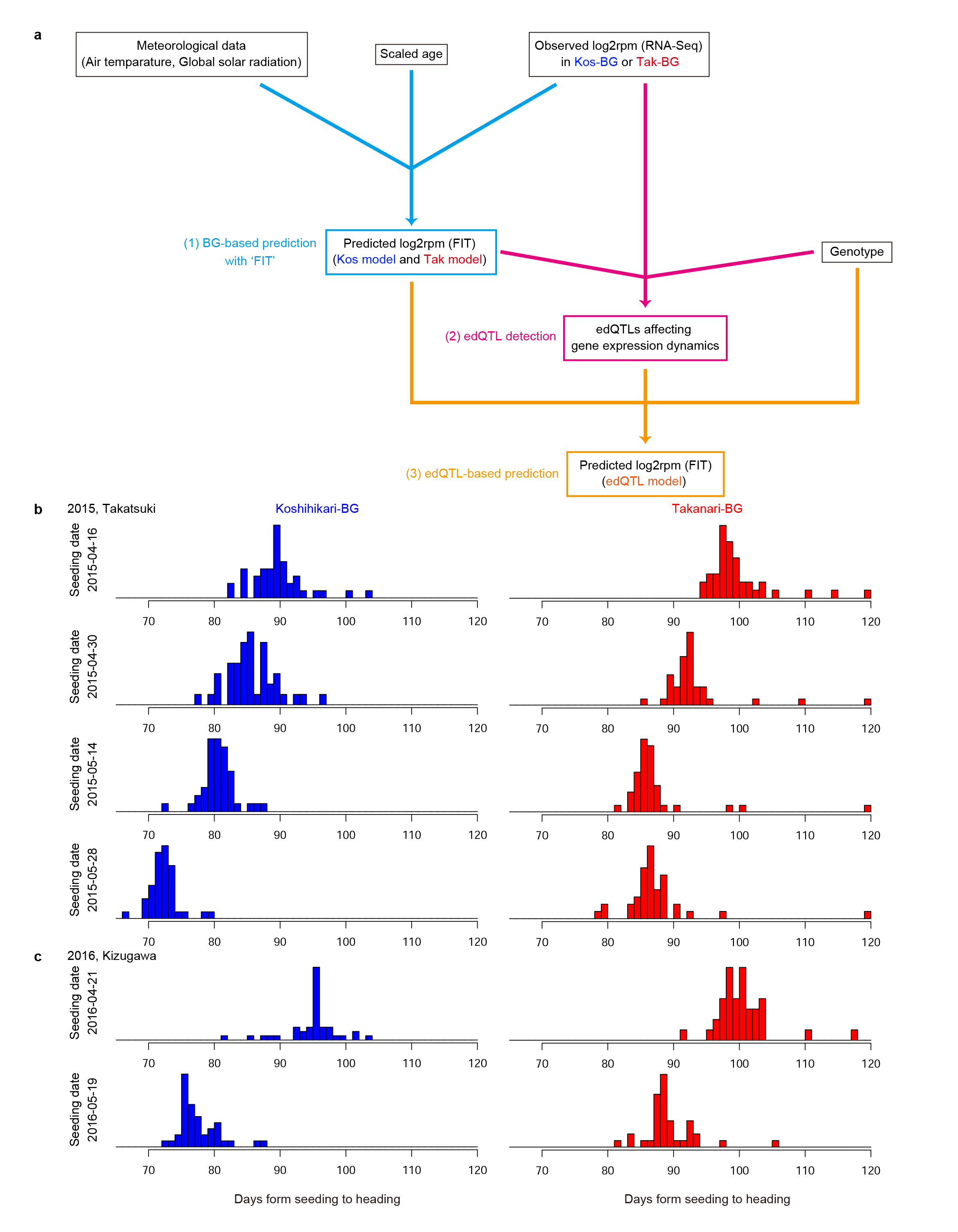

**Supplementary Figure 5 | Construction of the predictive model considering environmental factors and genotype. a,** Workflow of the analysis to identify and validate expression dynamics quantitative trait loci (edQTL). First, prediction models for ‘Koshihikari’ (Kos) and ‘Takanari’ (Tak) were developed with R package ‘FIT’, based on gene expression data in ‘Koshihikari’ background-genotype (Kos-BG) and Tak-BG samples and other factors. Second, edQTL were detected. Third, detected edQTL were validated by transcriptome prediction considering edQTL. **b,c,** Average length from seeding to heading of each ‘Koshihikari’ BG and ‘Takanari’ BG line in each transplant set in 2015 (**b**) and 2016 (**c**). Blue and red indicate Kos-BG and Tak-BG lines, respectively.

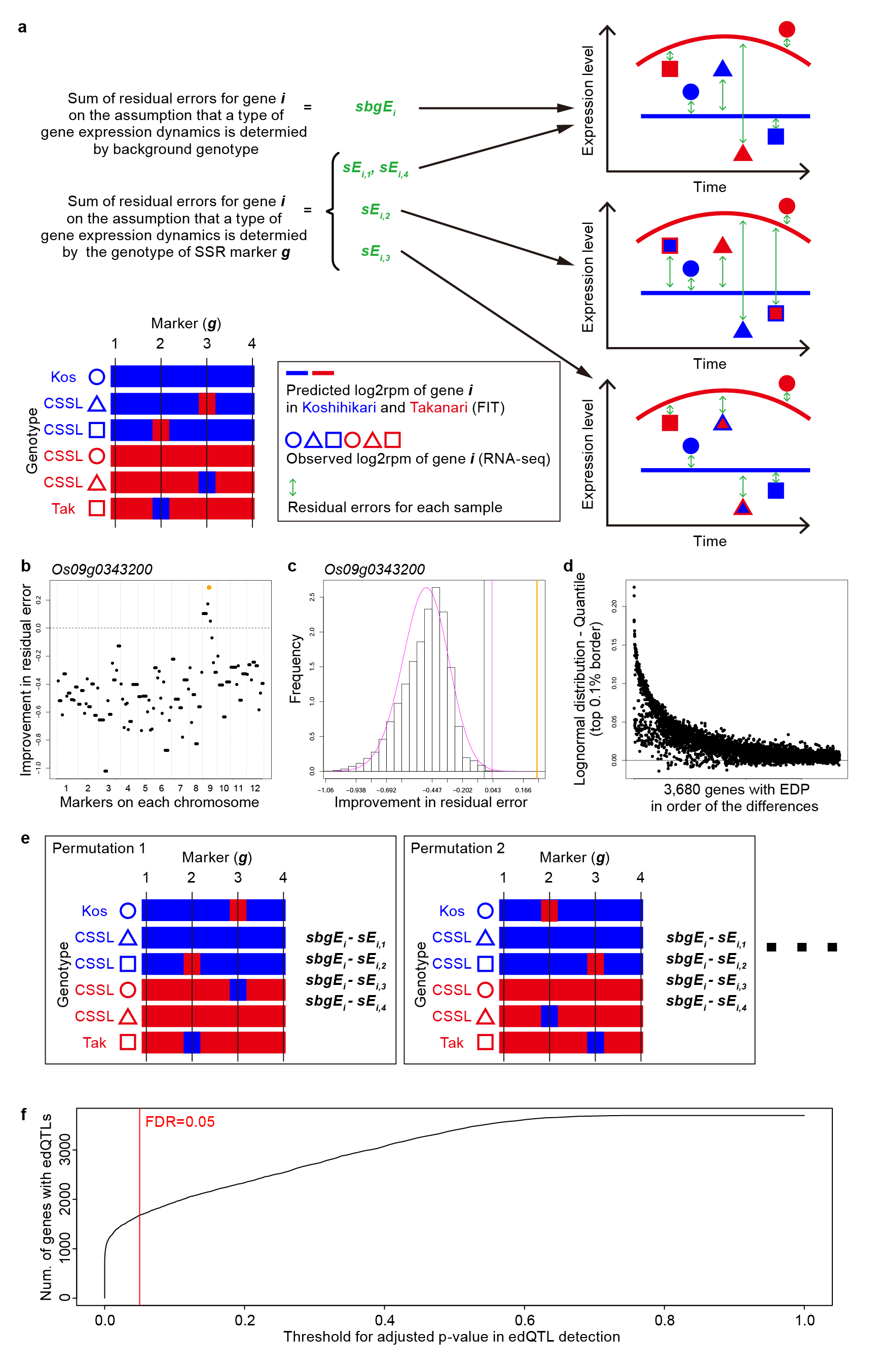
**Supplementary Figure 6 | Statistical test for expression dynamics quantitative trait loci (edQTL) detection. a,** The association between genetic variations and gene expression polymorphisms was evaluated by calculating residual errors in gene expression prediction. It was assumed that the type of each gene expression dynamics is determined by the background-genotype or SSR marker. In the example, edQTL affecting gene ***i*** exist around SSR marker *3*. In this case, the sum of residual errors ***sE_i,3_*** is smaller than the other residual errors (***sbgE_i_, sE_i,1_, sE_i,2_*** and ***sE_i,4_***). **b,** Improvement in the sum of residual errors for *Os09g0343200* (***sbgE_Os09g0343200_ - sE_Os09g0343200,i_***). The X-axis represents markers in order of chromosomal position. The dashed vertical lines indicate the border of each chromosome. The orange point indicates a marker of best improvement in sums of residual errors by the assumption that edQTL exist around each SSR marker. **c,** Distribution of the improvement in the sums of residual errors for *Os09g0343200* generated by permutation. The black line indicates the top 0.1% border in the distribution. Lognormal distributions were fitted to the distribution to calculate *p*-values. The magenta curve indicates the fitted lognormal distribution. The magenta vertical line represents the top 0.1% border of the lognormal distribution. Orange represents the degree of the best improvement in the residual error by the assumption that edQTL exists around each SSR marker. **d,** Differences between the top 0.1% border of the distributions generated by the permutation and the top 0.1% border of the fitted lognormal distributions of genes with expression dynamics polymorphism (EDP). **e,** Permutation of genotypes among ‘Koshihikari’- and ‘Takanari’-background lines; 1000 permutations were performed for statistical evaluation of the improvement in sum of residual errors by the assumption of edQTL. **f,** The relationship between the threshold for adjusted *p*-value in edQTL detection and the number of genes with edQTL.

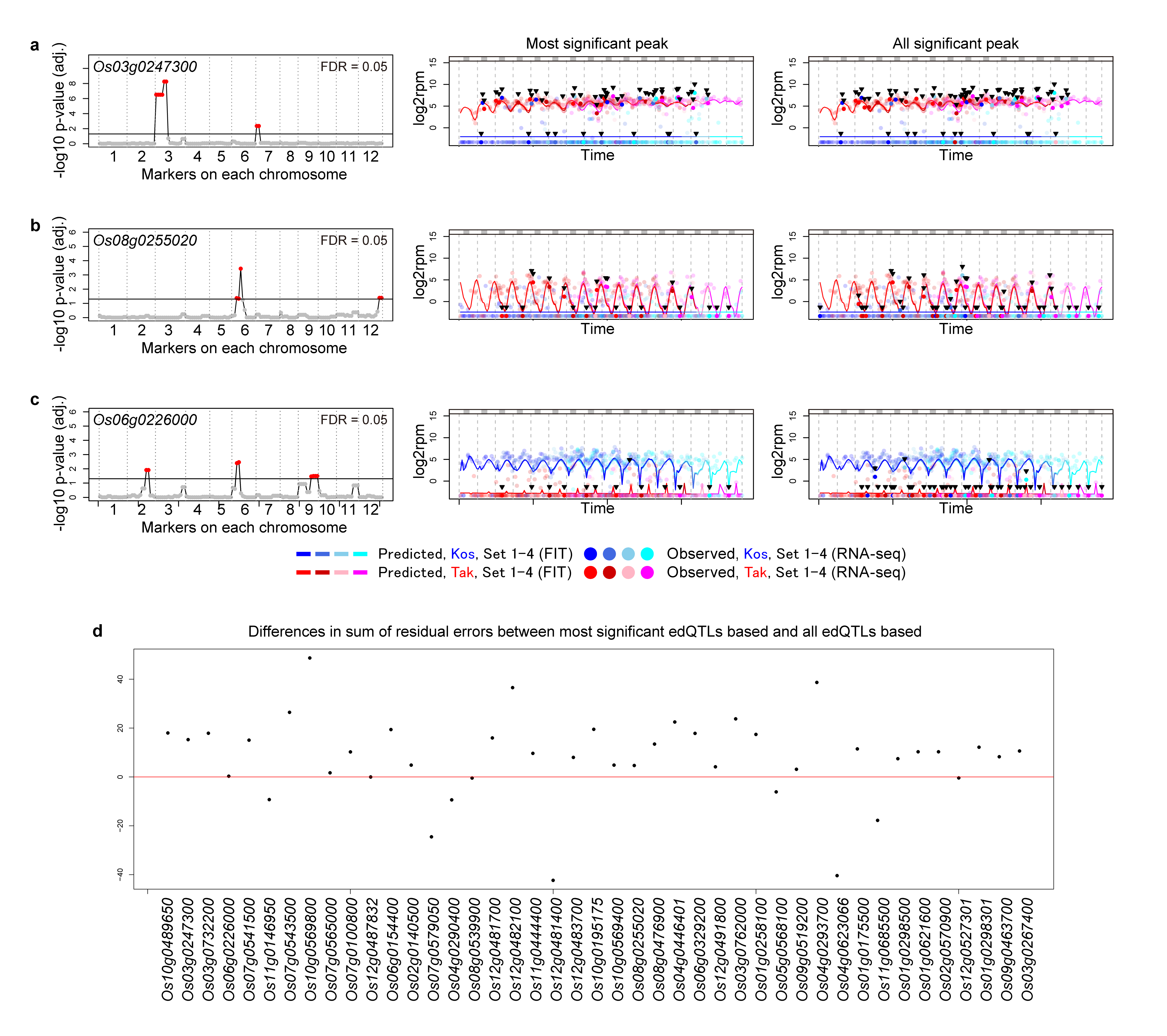
**Supplementary Figure 7 | Genes regulated by multiple expression dynamics quantitative trait loci (edQTL). a-c**, Examples of genes affected by multiple edQTL. [left panels] The result of edQTL detection. The X-axis represents markers in order of chromosomal position. [middle and right panels] Prediction of expression levels in Takatsuki in 2015 based on environmental information (time, air temperature, and solar radiation). Blue and red/pink lines indicate predicted expression levels in ‘Koshihikari’ and ‘Takanari’ in Transplant sets 1, 2, 3, and 4, respectively. Blue and red/pink points indicate the observed expression levels of samples in Transplant sets 1, 2, 3, and 4 of individuals with ‘Koshihikari’- and ‘Takanari’-type edQTL for the most significant peak [middle panels] or for all significant peaks [right panels], respectively. Points in intense colors emphasized by the arrowheads indicate samples harboring edQTL different from their background genotypes and light colors indicate samples harboring edQTL identical to their background genotypes. The upper gray bars indicate dark periods (global solar radiation < 0.3 kJ m^-2^ min^-1^). (**a**) *Os03g0247300*, (**b**) *Os08g0255020*, and (**c**) *Os06g0226000*. **d**, Degree of improvement in the sum of residual errors by considering all edQTL compared with that considering the most significant edQTL for the genes affected by several edQTL.

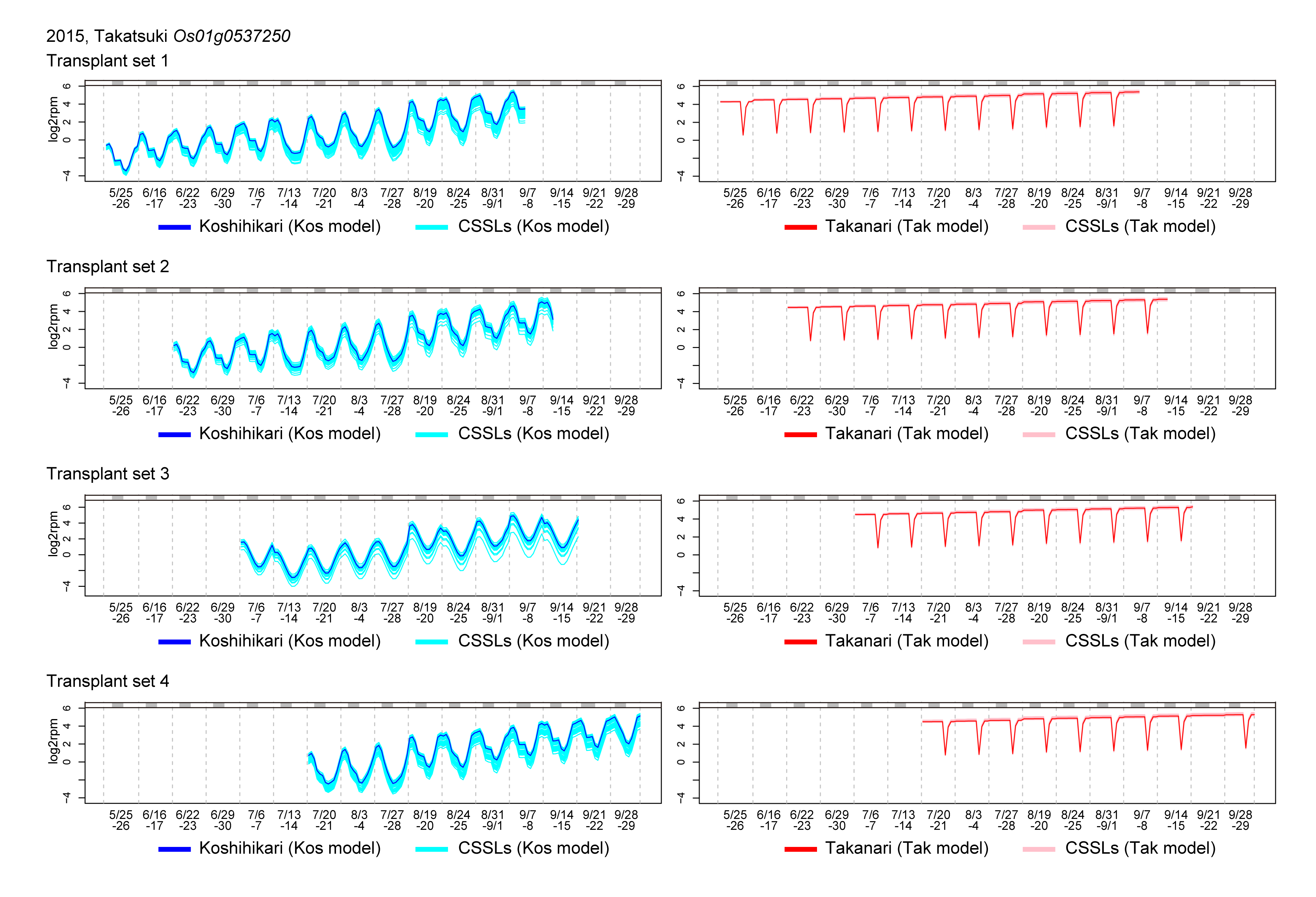

**Supplementary Figure 8 | Scaled age affected the prediction of gene expression dynamics.** An example (*Os01g0537250*) of the effect of scaled age on the prediction of expression dynamics in Takatsuki in 2015 in each transplant set. Blue and red/pink lines indicate predicted expression levels in ‘Koshihikari’ and ‘Koshihikari’-background-genotype CSSLs and ‘Takanari’ and ‘Takanari’-background-genotype CSSLs in Transplant sets 1, 2, 3, and 4, respectively. The upper gray bars indicate dark periods (global solar radiation < 0.3 kJ m^-2^ min^-1^).

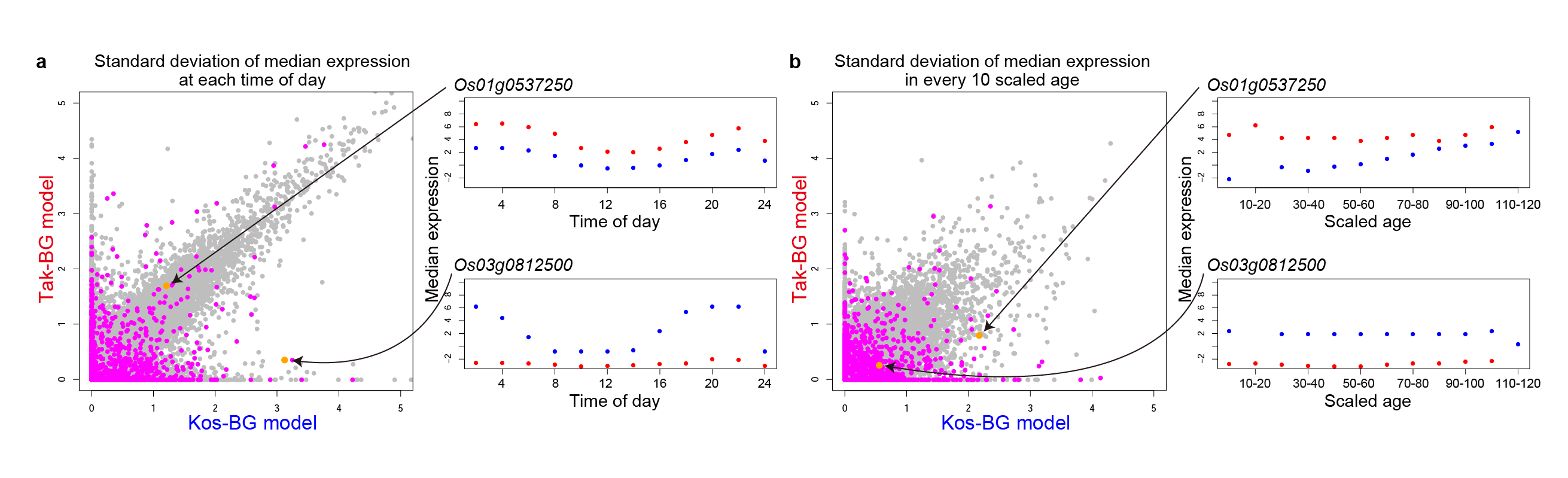

**Supplementary Figure 9 | Detection of expression dynamics quantitative trait loci (edQTL) related with differences in fluctuating environmental responses between ‘Koshihikari’ (Kos) and ‘Takanari’ (Tak). a,** Standard deviation of median gene expression levels at each time of day were shown. The standard deviation represents the degree of fluctuation of each gene expression with time of day. **b**, Standard deviation of median gene expression levels at each plants’ scaled age are shown. The standard deviation represents the degree of fluctuation of each gene expression with plants’ age. The genes depending on the time of day were defined as genes whose standard deviation of median expression at each time of day in either parental model was > 1. The genes depending on scaled age were defined as genes whose standard deviation of median expression in every 10 scaled age interval in either parental model was > 1. Magenta indicates genes regulated by edQTL. For example, the expression of *Os01g0537250* fluctuated with time of day and scaled age in both the Kos- and Tak-background -genotype (BG) models. In contrast, the expression of *Os03g0812500* fluctuated with time only in the Kos-BG model.

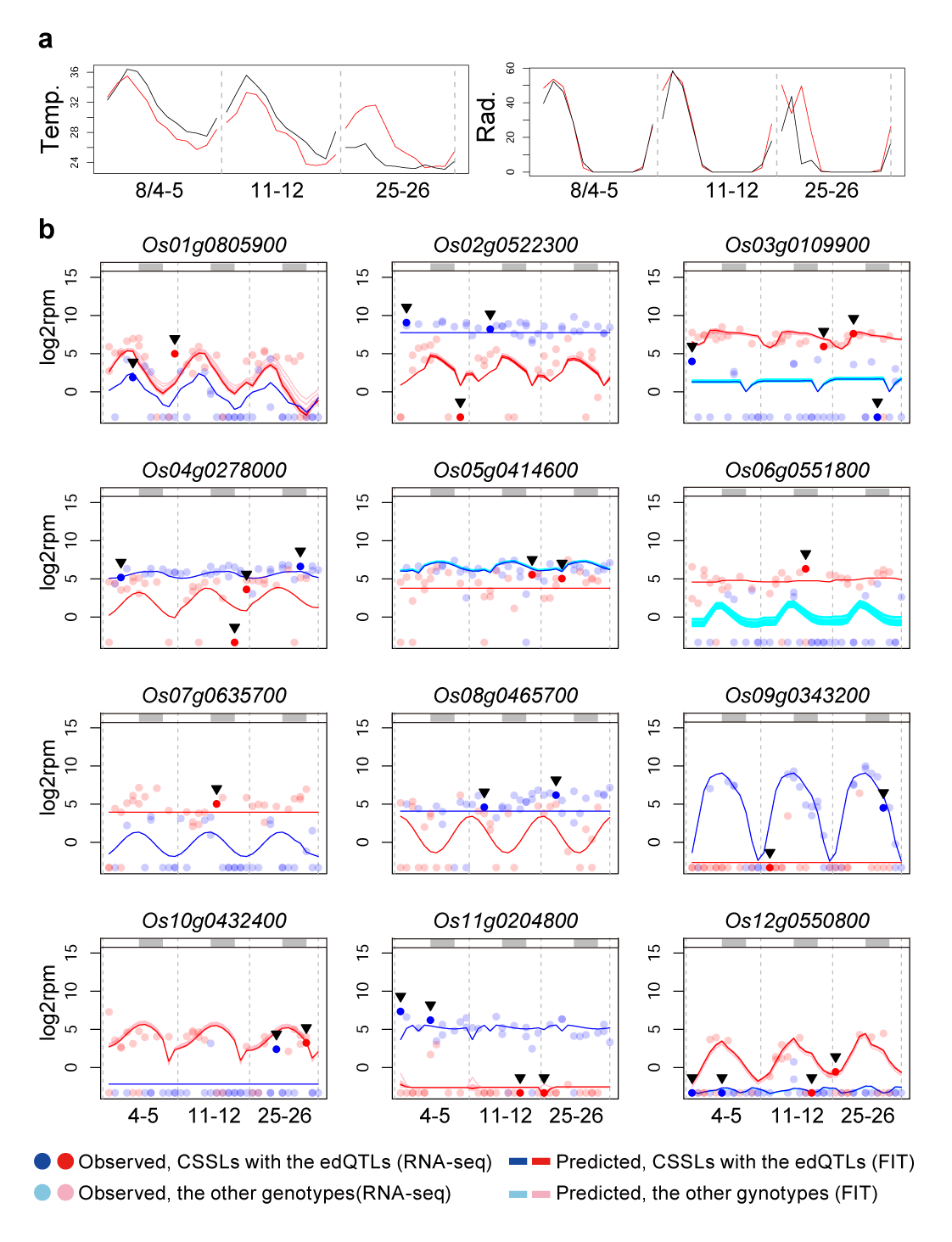

**Supplementary Figure 10 | Examples of prediction for expression dynamics in Kizugawa in 2016 based on environmental information and expression dynamics quantitative trait loci (edQTL).** **a,** Plots of meteorological data [air temperature (Temp., °C) and global solar radiation (Rad., kJ m^-2^ min^-1^)] on sampling dates in August. The black line indicates data in Takatsuki in 2015 and the red line indicates data in Kizugawa in 2016. **b,** Examples of observed and predicted gene expressions. Points in intense colors emphasized by arrowheads indicate samples harboring edQTL different from their background genotypes and light colors indicate samples harboring edQTL identical to their background genotypes in Transplant set 2. The upper gray bars indicate dark periods (global solar radiation < 0.3 kJ m^-2^ min^-1^).

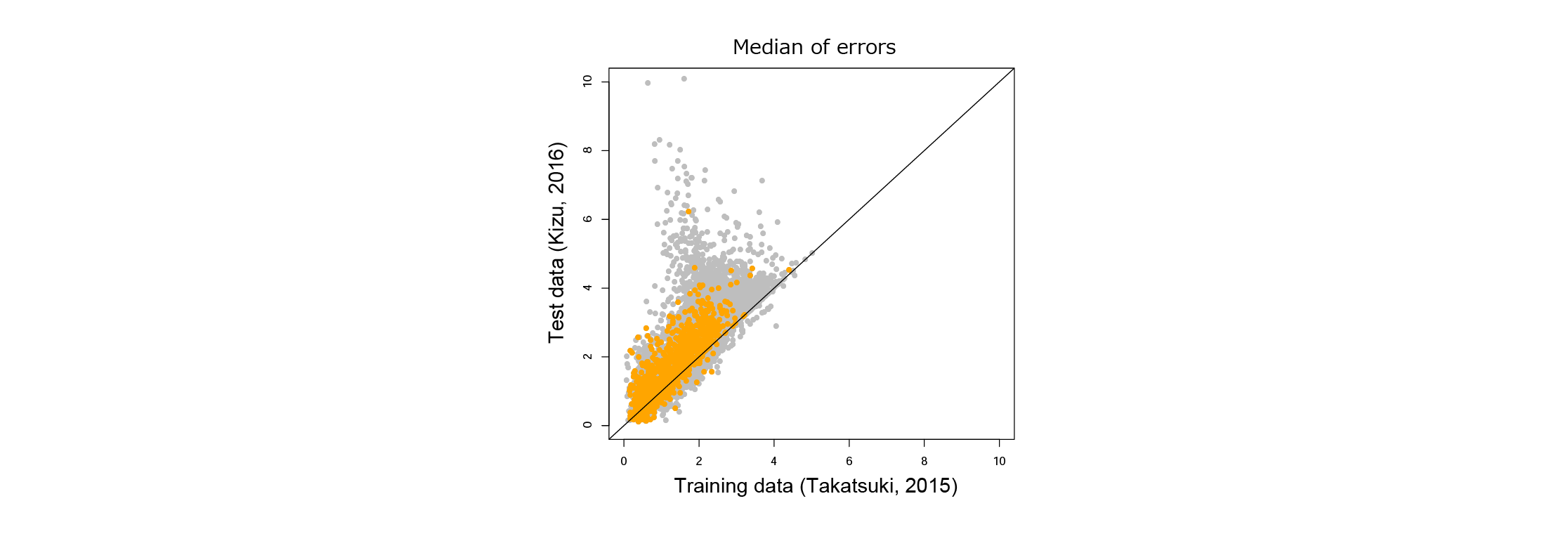

**Supplementary Figure 11 | Residual errors of gene expression by the expression dynamics quantitative trait loci (edQTL) model in 2015 (training data) and prediction errors in 2016.** Orange points indicate genes that were affected by edQTL and gray points indicate genes that were not affected by edQTL.

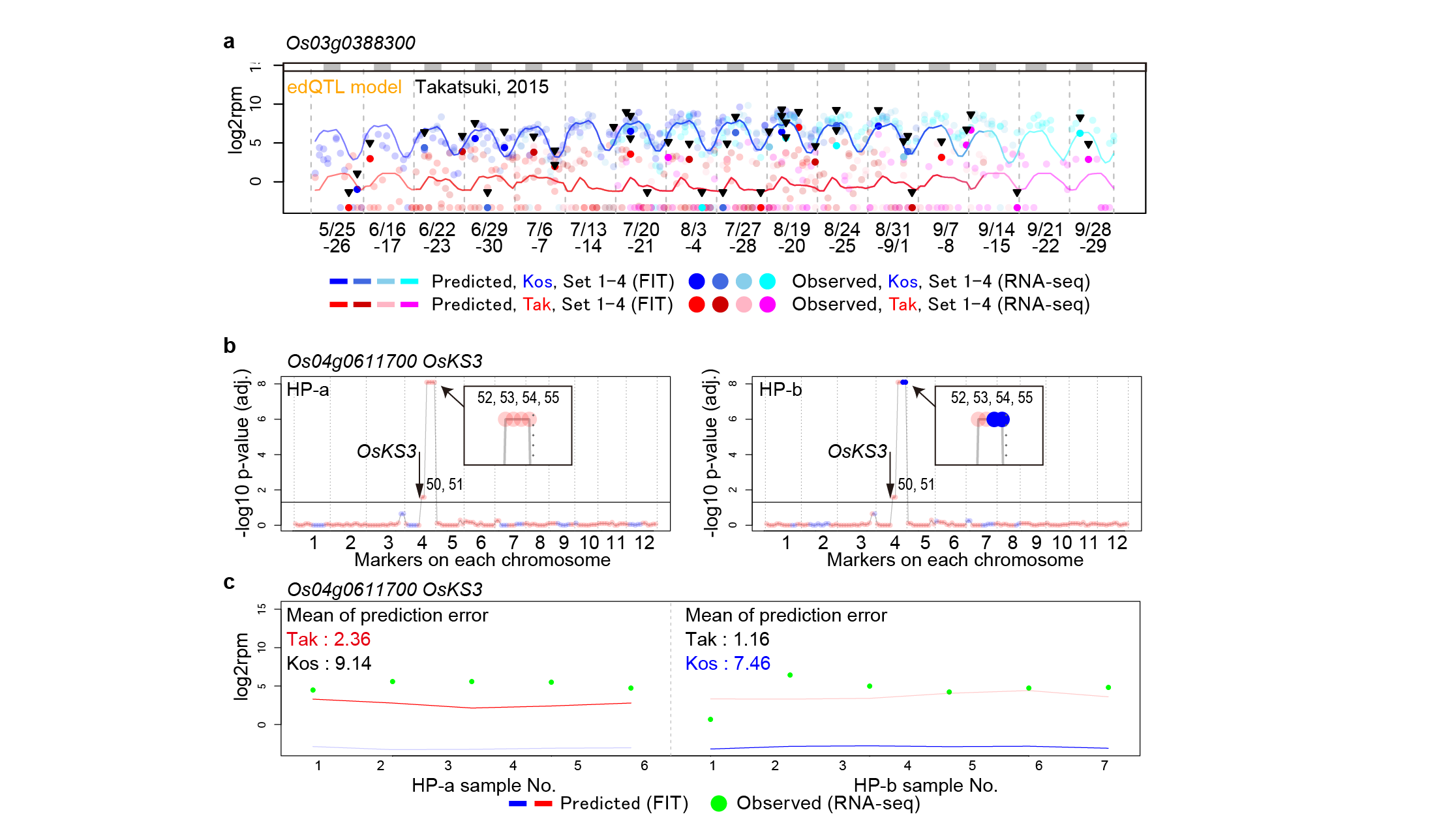

**Supplementary Figure 12 | Limitation of expression dynamics quantitative trait loci (edQTL) detection using chromosome segment substitution lines (CSSLs). a,** Prediction of *Os03g0388300* expression level in ‘Koshihikari’ and ‘Takatsuki’ in 2015 based on scaled age and environmental information (time, air temperature, and solar radiation). Blue and red/pink lines indicate predicted expression levels in ‘Koshihikari’ and ‘Takanari’ in Transplant sets 1, 2, 3, and 4, respectively. Blue and red/pink points indicate observed expression levels of samples in Transplant sets 1, 2, 3, and 4 of individuals with ‘Koshihikari’-type edQTL and ‘Takanari’-type edQTL, respectively. Intense colors emphasized by arrowheads indicate samples harboring edQTL different from their background genotypes and light colors indicate samples harboring edQTL identical to their background genotypes. The upper gray bars indicate dark periods (global solar radiation < 0.3 kJ m^-2^ min^-1^). **b,** edQTL for *OsKS3* and genotypes of HP-a and HP-b. The red and blue points indicate ‘Koshihikari’- and ‘Takanari’-type markers, respectively. Dark blue points indicate significant ‘Koshihikari’-type edQTL. As a result of edQTL detection of *OsKS3*, four markers were identified as edQTL affecting the expression dynamics. The two markers (No. 54 and 55) are substituted with ‘Koshihikari’-type in HP-b. **c,** Prediction of *OsKS3* expression in HP-a and HP-b. Strongly colored lines are applied models for HP-a and HP-b.

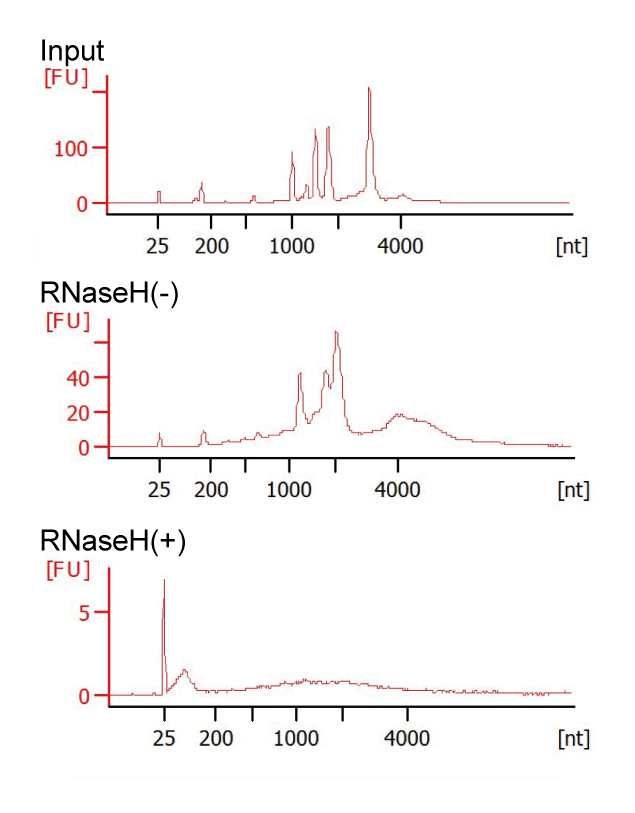

**Supplementary Figure 13 | Selective-depletion of abundant transcripts.** RNA was analyzed with a Bioanalyzer and Agilent RNA 6000 Nano Assay. Selective-depletion using antisense oligo and RNase H was conducted. After enzymatic degradation, peaks of highly abundant transcripts disappeared.

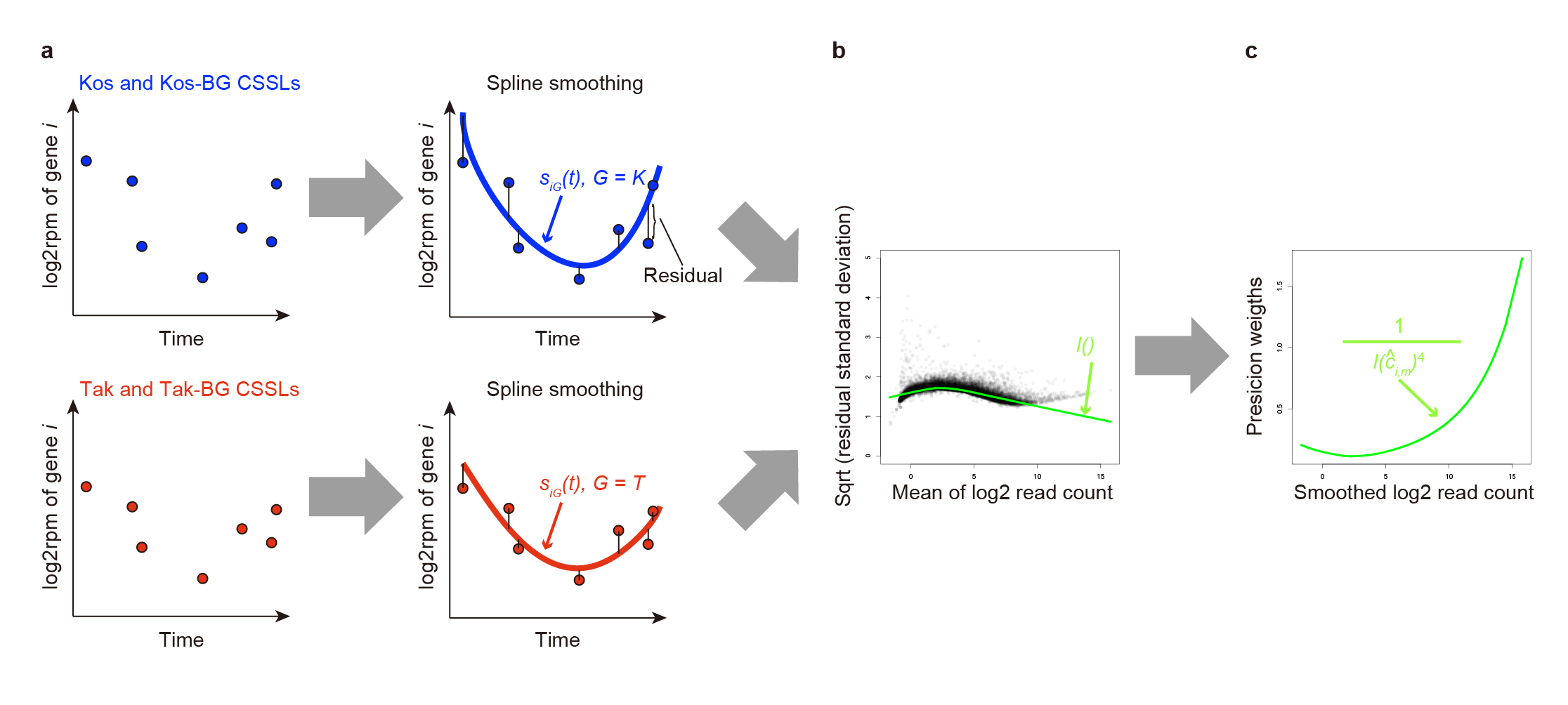

**Supplementary Figure 14 | Calculation of the precision weights matrix. a,** Spline smoothing was conducted for time-series of observed log2rpm values of each gene in samples of ‘Koshihikari’- or ‘Takanari’-background genotypes, resulting in function values ***s_i,G_(t)***. Then, residual errors between smoothed log2rpm and observed log2rpm were calculated. **b,** Fitting of the LOWESS curve to the square-root of the residual standard deviations as a function of mean log2count yielded a function value ***l()***. **c,** Precision weights for observed log2rpm were calculated based on smoothed log2 read count of each gene in each sample.

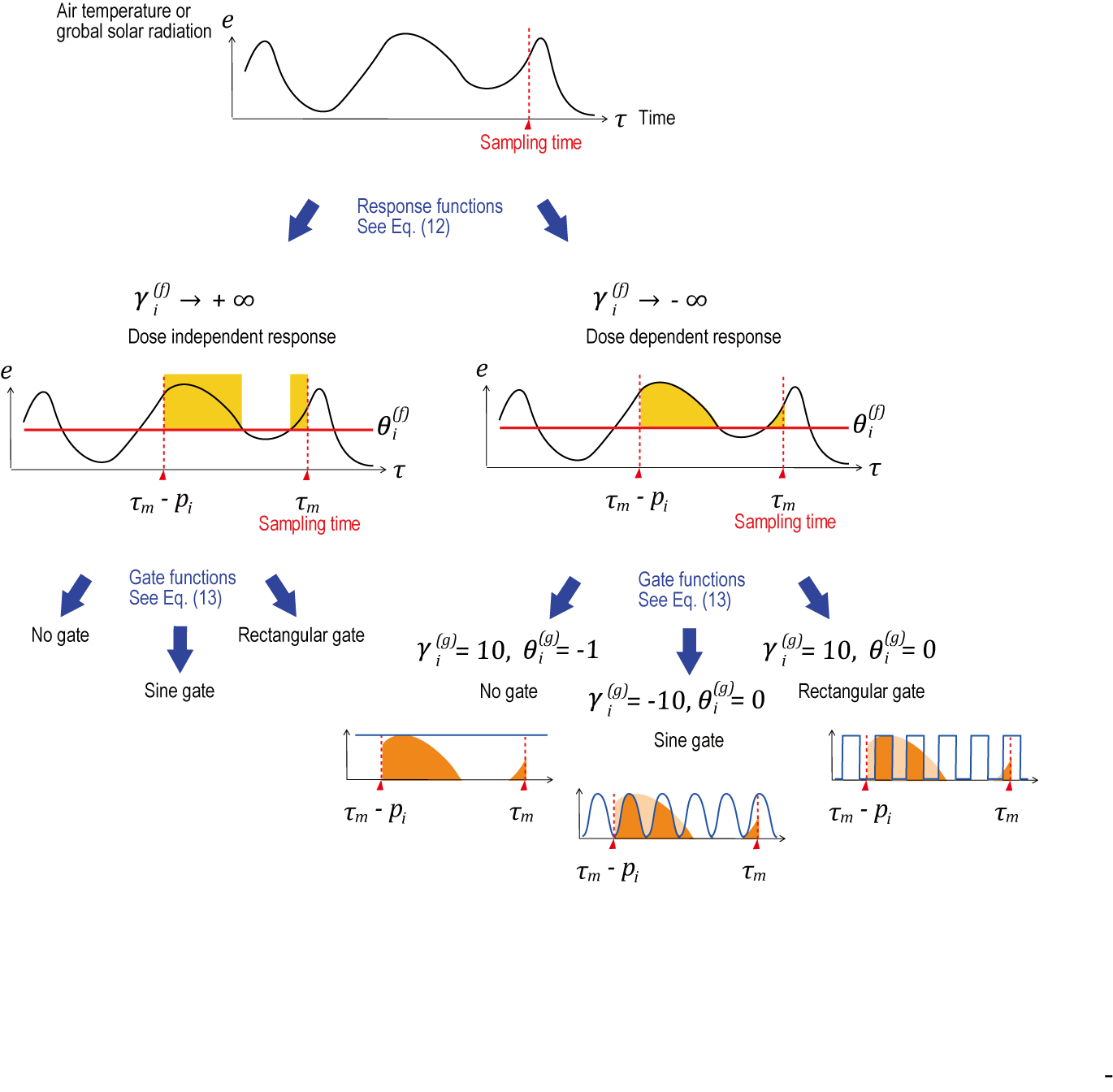

**Supplementary Figure 15 | Schematic figures of the calculation of gene expression response to environmental stimuli.** The top plot displays meteorological data (vertical axis, *e*) and sampling time (horizontal axis, *τ*). The middle plots indicate typical examples of response modes. The yellow area indicates response amplitude before diurnal gating. The red line represents threshold, *𝜃_i_^(f)^*. *p_i_* indicates a period in which an environmental stimulus affects the expression. Bottom plots indicate typical examples of diurnal gates. The orange area indicates response amplitude after diurnal gating. The blue line represents diurnal gate.

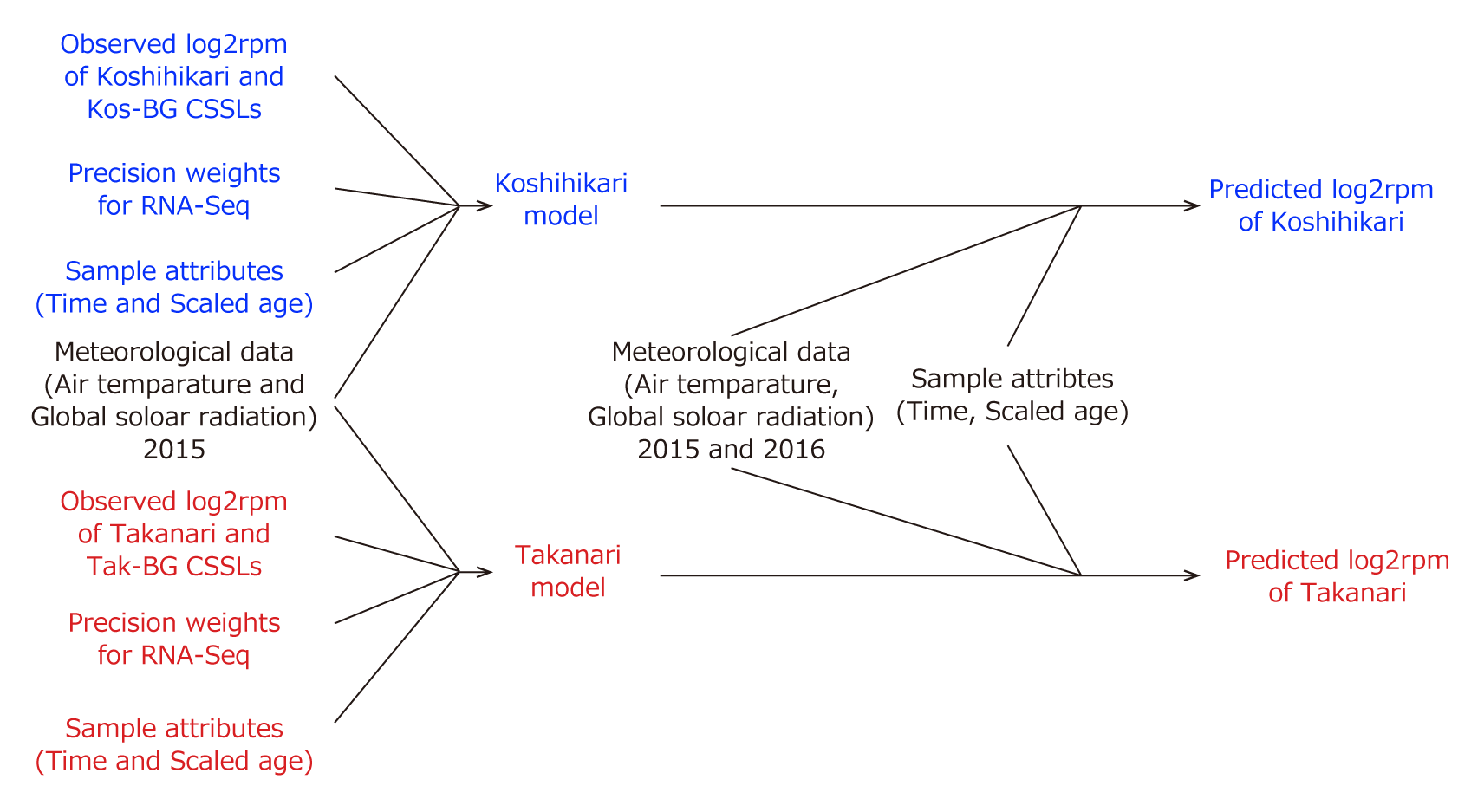

**Supplementary Figure 16 | Overview of model development and prediction of gene expression with “FIT”**. Predictive models of gene expression dynamics for ‘Koshihikari’ and ‘Takanari’ were developed with ‘FIT’ based on RNA-Seq data (observed log2rpm), the corresponding precision weights, meteorological data, and scaled age. Then, based on the models with input of meteorological data and sample attributes, predicted log2rpm of ‘Koshihikari’ and ‘Takanari’ can be obtained.
